# Reactivation of chromosome signalling induces reversal of the meiotic program

**DOI:** 10.1101/754341

**Authors:** Maikel Castellano-Pozo, Sarai Pacheco, Georgios Sioutas, Angel Luis Jaso-Tamame, Marian H Dore, Enrique Martinez-Perez

## Abstract

Chromosome movements and programmed DNA double-strand breaks (DSBs) promote homologue pairing and initiate recombination at meiosis onset. Meiotic progression involves checkpoint-controlled termination of these events when all homologue pairs achieve synapsis and form crossover precursors. We show that termination of chromosome movement and DSB formation is reversible and is continuously implemented by the synaptonemal complex (SC), which silences chromosome signals that promote CHK-2 activity. Forced removal of the SC or different meiosis-specific cohesin complexes, which are individually required for SC stability, causes rapid CHK-2-dependent reinstallation of the DSB-formation and chromosome-movement machinery. This nuclear reorganization occurs without transcriptional changes, but requires signalling from HORMA protein HTP-1. Conversely, CHK-2 inactivation causes rapid disassembly of the DSB-formation and chromosome-movement machinery. Thus, nuclear organization is constantly controlled by the level of CHK-2 activity. Our results uncover an unexpected plasticity of the meiotic program and show how chromosome signalling integrates nuclear organization with meiotic progression.

## Introduction

The formation of haploid gametes from diploid germ cells during meiosis constitutes a cornerstone of sexual reproduction. Defects in this process cause sterility and aneuploid gametes that impair fitness of the resulting progeny. Accurate transmission of chromosomes into the gametes depends critically on the establishment of crossover (CO) events between paternal and maternal homologous chromosomes (homologues) during the long prophase preceding the first meiotic division, when COs, together with sister chromatid cohesion (SCC), ensure correct chromosome orientation on the spindle (Petronczki et al., 2003). Thus, meiotic cells have evolved meiosis-specific chromosome structures and surveillance mechanisms to ensure that every pair of homologues is connected by COs before nuclei proceed to the first meiotic division.

Meiotic chromosome morphogenesis requires the assembly of proteinaceous axial elements containing meiosis-specific versions of cohesin, the complex that provides SCC. In yeast loading of cohesin containing Rec8, a meiosis-specific version of cohesin’s mitotic kleisin Scc1 (Klein et al., 1999), ensures proper meiotic chromosome function. Higher eukaryotes use additional meiosis-specific kleisins, including Rad21L in mouse and the highly identical COH-3 and COH-4 in *C. elegans* (Ishiguro, 2019; Pasierbek et al., 2001; Severson et al., 2009). In addition to establishing SCC, loading of meiotic cohesin promotes recruitment of HORMA-domain proteins (HORMADs) to axial elements, rendering chromosomes competent to initiate meiotic recombination, via the formation of DNA double strand breaks (DSBs), and to undergo pairing of homologous chromosomes. Homologue pairing is facilitated by cytoskeleton-driven chromosome movements during early meiotic prophase and culminates with the assembly of the synaptonemal complex (SC), a ladder-like structure that bridges together the axial elements of aligned homologues (Zickler and Kleckner, 2015). This process, known as synapsis, stabilises homologue interactions and is essential to ensure that a subset of DSBs become CO-designated sites during the pachytene stage of meiotic prophase, which is defined by full synapsis. Thus, the processes leading to CO formation are closely integrated with the establishment of meiosis-specific chromosome structures built over a cohesin scaffold. However, how different cohesin complexes and the SC contribute to meiotic chromosome function once full synapsis is achieved and CO precursors are formed remains poorly understood.

The formation and repair of DSBs into COs is mechanistically coupled to early meiotic progression by a network of surveillance mechanisms that monitor specific, but incompletely understood, pairing and recombination intermediates (Subramanian and Hochwagen, 2014). The presence (or absence) of these meiotic chromosome metabolism intermediates results in signals that feedback to regulate checkpoint kinases, which then target components of the pairing, recombination, and cell cycle machinery (Keeney et al., 2014). These quality control mechanisms fulfil two important roles. Firstly, they act to limit the temporal window during which nuclei remain competent for DSB formation, ensuring timely cessation of this activity once CO precursors are formed on all chromosomes, thus preventing the genotoxic effects of excess DSBs. Secondly, they regulate meiotic progression in a checkpoint manner, inducing arrest of nuclei at the stage when defects in specific CO-promoting events are first detected. For example, mutations that impair DSB processing into CO precursors cause extension of DSB-permissive stages in yeast and *C. elegans* (Rosu et al., 2013; Stamper et al., 2013; Thacker et al., 2014). The regulated exit from DSB-permissive stages represents a fundamental transition of the meiotic program, but how nucleus-wide loss in competence for DSB formation is sustained remains unclear.

The *C. elegans* germline provides a powerful system to investigate how feedback mechanisms integrate pairing and recombination with early meiotic progression (Hillers et al., 2017). CHK-2 promotes DSB formation, chromosome movement, and SC assembly during early prophase and the temporal window of CHK-2 activity is controlled by feedback from the progression of these events mediated by HORMADs (Kim et al., 2015; MacQueen and Villeneuve, 2001; Martinez-Perez and Villeneuve, 2005). Despite sharing some components, these feedback mechanisms are mechanistically different from checkpoints that induce apoptosis of nuclei with persistent DNA damage or asynapsed chromosomes at late pachytene (Bhalla and Dernburg, 2005; Gartner et al., 2000). Importantly, SC assembly is independent of recombination in worms (Dernburg et al., 1998) and both processes are under surveillance to feedback on CHK-2 (Woglar et al., 2013). In wild-type germlines synapsis induces termination of CHK-2-depedent chromosome movements by pachytene entrance, while the formation of CO precursors triggers loss of CHK-2-dependent markers of DSB formation by mid pachytene. Once achieved, nucleus-wide loss of CHK-2 activity is thought of as a unidirectional transition of the meiotic program that leads to the completion of recombination and progression towards chromosome segregation.

In the current work, we combine the experimental advantages of the *C. elegans* germline with temporally-resolved protein removal methods to investigate how cohesin and the SC contribute to meiotic chromosome function once full synapsis and early recombination steps are completed. We uncover a role for REC-8 and COH-3/4 cohesin in promoting SC stability and demonstrate that direct or indirect, via cohesin removal, SC disassembly triggers rapid and nucleus-wide redeployment of the pairing and recombination machinery, inducing drastic changes in nuclear organization. This apparent reversal in meiotic progression requires CHK-2 reactivation mediated by HORMA protein HTP-1, but occurs in the absence of transcriptional changes. Our findings have important implications for understanding the quality control mechanisms that ensure fertility in higher eukaryotes.

## Results

### Time-resolved removal of specific cohesin complexes from pachytene nuclei

*C. elegans* expresses two types of meiosis-specific cohesin complexes defined by their kleisin subunit: REC-8 and the highly identical and functionally redundant COH-3 and COH-4 (referred to as COH-3/4 from now on). Both REC-8 and COH-3/4 are prominent components of pachytene axial elements and mutant analysis has uncovered overlapping roles for REC-8 and COH-3/4 cohesin in axis assembly, as well as non-overlapping roles in SCC, pairing, synapsis, and recombination (Crawley et al., 2016; Pasierbek et al., 2001; Severson et al., 2009; Severson and Meyer, 2014). Crucially, however, *rec-8* and *coh-3/4* mutants display severe chromosome organization defects from the onset of meiosis, making mutant analysis impractical for clarifying the roles of REC-8 and COH-3/4 complexes during pachytene. To bypass this intrinsic limitation of kleisin mutant analysis, we created kleisin versions that can be removed from meiotic chromosomes in a temporally-controlled manner by introducing 3 repeats of the TEV protease recognition motif in REC-8 (after Q289) and COH-3 (after 1315). TEV-mediated cleavage of kleisin subunits mimics cleavage by separase at anaphase onset, and this approach has been successfully used to rapidly remove cohesin from chromosomes *in vivo* in other organisms (Oliveira et al., 2010; Tachibana-Konwalski et al., 2010). Both REC-8^3XTEV^::GFP and COH-3^3XTEV^::mCherry were fully functional, as single copy transgenes expressing REC-8 and COH-3 from their respective promoter and 3’ UTRs complemented the meiotic defects of *rec-8* and *coh-3 coh-4* mutants respectively (Figures S1A-B). Unless otherwise indicated, TEV experiments described below were performed in germlines from worms expressing REC-8^3XTEV^::GFP or COH-3^3XTEV^::mCherry in backgrounds carrying null mutations in *rec-8* or *coh-3 coh-4* respectively.

The sequential stages of meiotic prophase are easily identified in the *C. elegans* germline based on the position of nuclei within the germline, nuclear appearance, and by cytological markers of specific meiotic events (Figure 1A). Nuclei in leptotene-zygotene stages (transition zone) are characterised by chromosome clustering and by markers of chromosome movement, SC assembly, and DSB formation. Entry into pachytene is marked by chromosome dispersal, high numbers of early recombination intermediates, and full SC assembly. By late pachytene, markers of early recombination disappear and CO-designated sites emerge. Since germ cells in *C. elegans* are syncytial, we reasoned that microinjection of the TEV protease into the germline of live worms would allow us to remove REC-8^3XTEV^::GFP or COH-3^3XTEV^::mCherry from the large population of nuclei at the pachytene stage. Indeed, germlines dissected and fixed 3.5 hours after TEV injection demonstrated efficient removal of REC-8^3XTEV^::GFP or COH-3^3XTEV^::mCherry from axial elements at all stages of meiotic prophase while non-targeted complexes remained bound to chromosomes (Figure 1B and Figures S1D-E). Importantly, as nuclei take about 36 hours to progress through pachytene (Jaramillo-Lambert et al., 2007) (Figure 1A), germline dissection 3.5 hours post-TEV injection allowed us to examine nuclei nearly at the same position that they occupied at the time of injection. TEV injection in control germlines expressing REC-8::GFP or COH-3::mCherry without the TEV motif confirmed that only TEV-tagged versions are removed following injection (Figure S1C). Thus, this TEV-based approach provides a powerful tool to rapidly and specifically remove different cohesin complexes from pachytene chromosomes.

**Figure 1.**
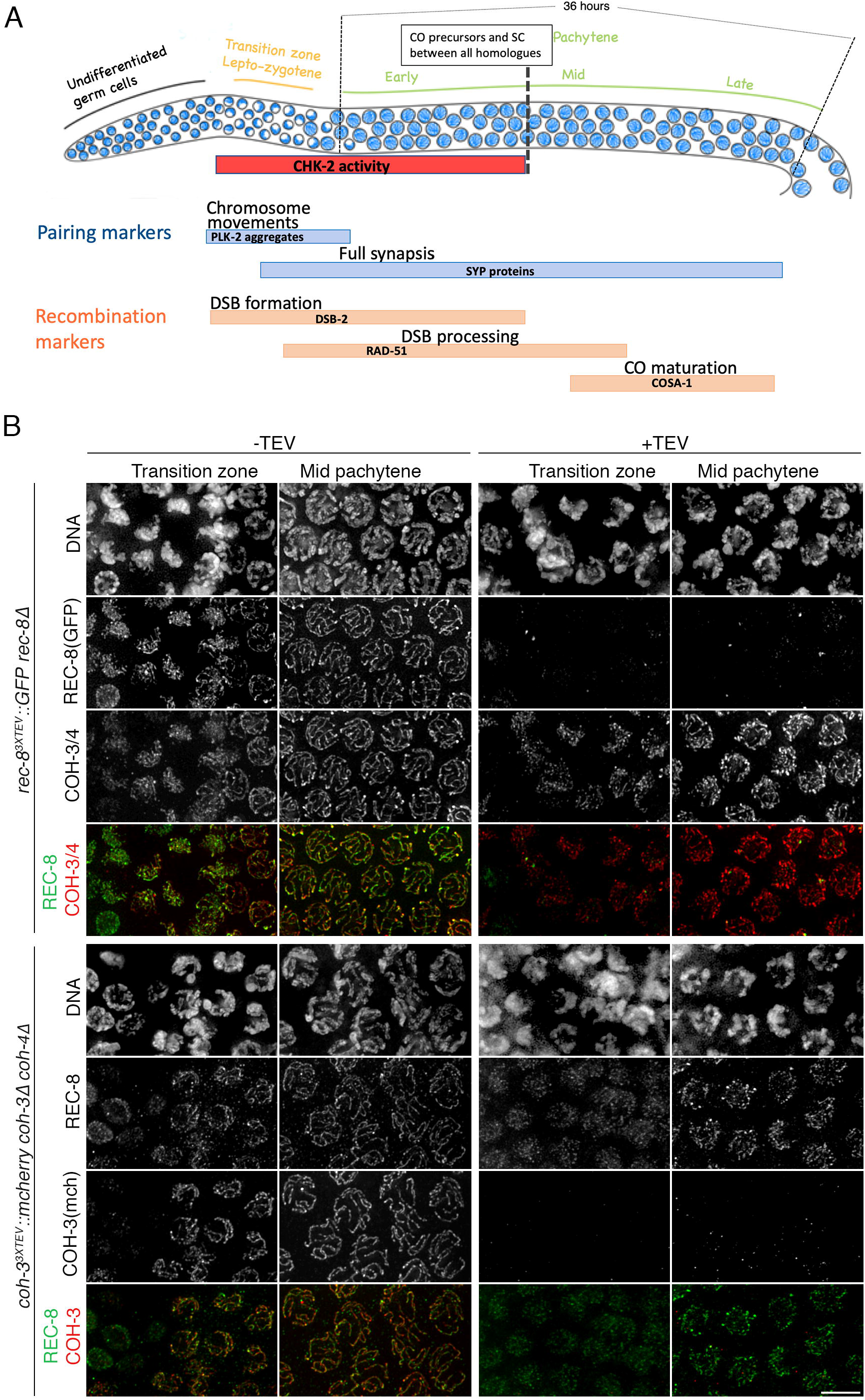
TEV-mediated cohesin removal in the *C. elegans* germline. **(A)** Diagram of a wild-type *C. elegans* germline indicating key meiotic events and the timing of appearance and disappearance of different cytological markers of pairing and recombination. **(B)** Projections of nuclei at the indicated stages before and after (3.5 hours) TEV injection demonstrating efficient and specific removal of REC-8^3XTEV^::GFP and COH-3^3XTEV^::mCherry. Scale bar= 5 μm. See also Figure S1.

### Removal of REC-8 or COH-3/4 cohesin induces SC disassembly

Having set up conditions to rapidly remove cohesin from chromosomes, we investigated the contribution of REC-8 and COH-3/4 complexes to axial elements and the SC in pachytene nuclei, as these structures are assembled over a cohesin scaffold during leptotene-zygotene. The central region of the SC in *C. elegans* is composed of four proteins (SYP-1-4) that are interdependent for their loading during early prophase (Colaiacovo et al., 2003; MacQueen et al., 2002; Smolikov et al., 2007; Smolikov et al., 2009). Using anti-SYP-1 antibodies we observed that removal of REC-8^3XTEV^::GFP or COH-3^3XTEV^::mCherry caused SC disassembly at all stages of meiotic prophase (Figure 2A and Figure S2A). In most germlines, REC-8^TEV^::GFP cleavage caused concentration of SYP-1 in a few aggregates, where it colocalized with remaining GFP (REC-8) signals (Figure 2A). These aggregates were more prominent in late pachytene nuclei, where they typically appeared as 6 chromosome-associated short stretches (Figure 2A). Since late pachytene nuclei in worms contain six crossover-designated sites, one per homologue pair, that are visible as 6 COSA-1 foci (Yokoo et al., 2012), we tagged the *cosa-1* gene with HA using CRISPR and crossed this allele into the REC-8^3XTEV^::GFP strain. Following TEV injection, we observed that each persistent REC-8^3XTEV^::GFP stretch in late pachytene was associated with a COSA-1 focus and most nuclei contained 6 COSA-1 foci, as did non-injected controls (Figure 2B). These results show that REC-8 and COH-3/4 are individually required for SC stability during pachytene and suggest that REC-8 cohesin is locally regulated around CO-designated sites.

**Figure 2.**
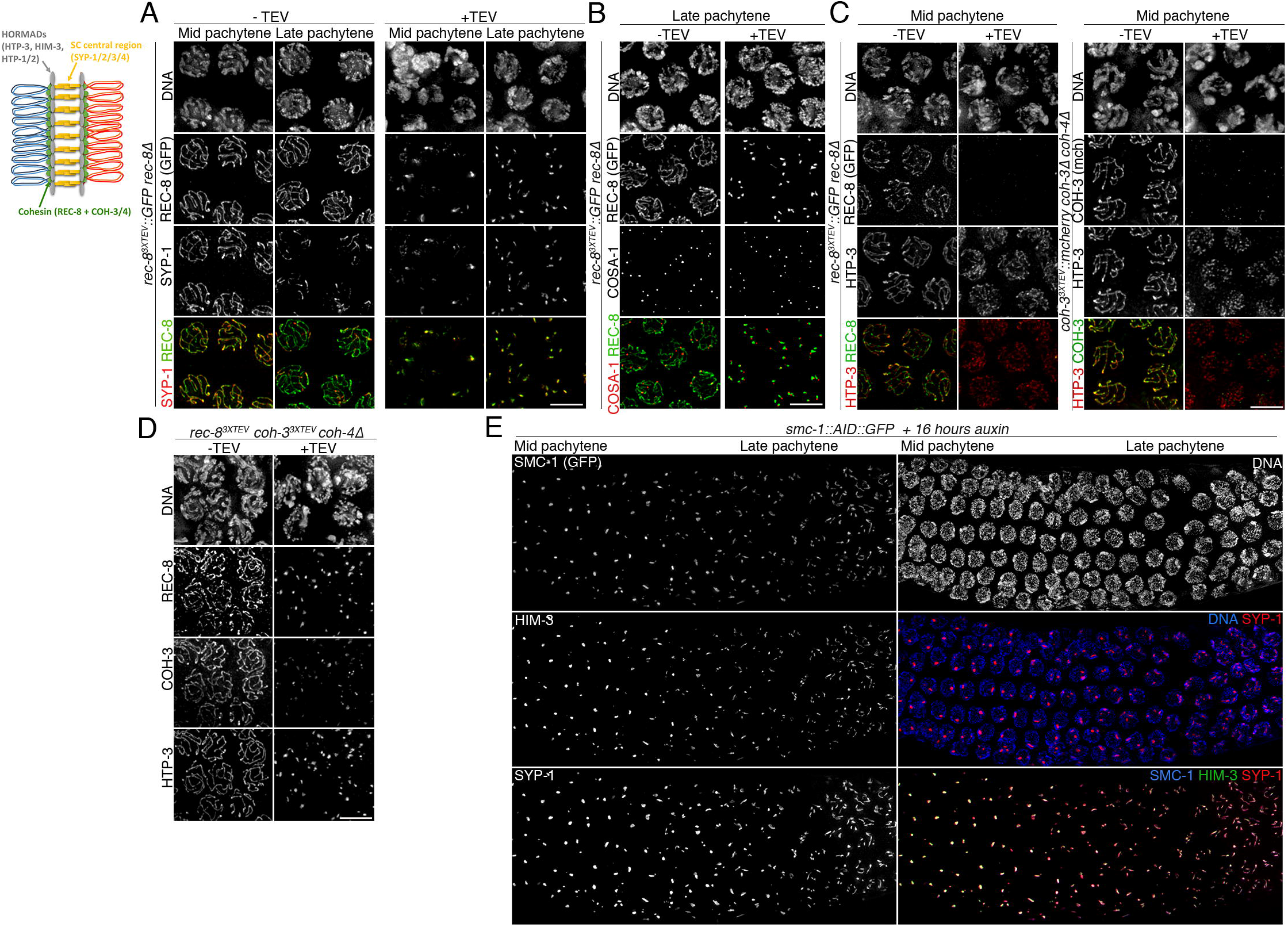
Cohesin promotes SC stability in pachytene nuclei. Diagram of a pair of synapsed homologues at the pachytene stage indicating the localization of axis and SC components. Panels A-E contain projections of nuclei at the indicated stages before and after (3.5 hours) TEV injection acquired on a Delta Vision microscope, while panel E shows nuclei after auxin degron-mediated depletion of SMC-1::AID:GFP acquired with a SIM microscope. **(A)** TEV-mediated removal of REC-8^3XTEV^::GFP induces SC disassembly. **(B)** COSA-1 foci persist following TEV-mediated removal of REC-8^3XTEV^::GFP. **(C)** Axial elements containing HTP-3 persist following TEV-mediated removal of REC-8^3XTEV^::GFP or COH-3^3XTEV^::mCherry. Note that HTP-3 signals appear more discontinuous when COH-3^3XTEV^::mCherry is removed. **(D)** Simultaneous depletion of REC-8^3XTEV^ and COH-3^3XTEV^ results in axial element disassembly. **(E)** Auxin degron-mediated depletion of SMC-1::AID::GFP for 16 hours induces axis and SC disassembly. Scale bar= 5 μm in all panels. See also Figure S2.

### Disassembly of axial elements requires simultaneous removal of REC-8 and COH-3/4

Axial elements in *C. elegans* require HORMA-domain protein HTP-3, which is recruited to chromosomes at meiotic onset in a cohesin-dependent manner (Goodyer et al., 2008; Lightfoot et al., 2011; Severson et al., 2009). HTP-3 then acts as a scaffold for the recruitment of additional HORMA proteins: HIM-3 and HTP-1/2, which play multiple roles in pairing, synapsis, recombination, and SCC release during the meiotic divisions (Couteau and Zetka, 2005; Ferrandiz et al., 2018; Kim et al., 2014; Zetka et al., 1999). All four HORMA proteins localise along the whole length of axial elements until late pachytene, when crossovers trigger restricted removal of HTP-1/2 (Martinez-Perez et al., 2008). In contrast to SC central region components, all HORMA proteins remained associated with axial elements following cleavage of REC-8^3XTEV^::GFP or COH-3^3XTEV^::mCherry (Figures 2C and Figures S2B-C). We noted that staining of HORMA proteins became weaker and more discontinuous following removal of COH-3^3XTEV^::mCherry than REC-8^3XTEV^::GFP, suggesting that COH-3/4 cohesin makes a larger contribution to axis integrity in pachytene nuclei, in agreement with the weaker axis appearance observed in *coh-3 coh-4* double mutants when compared to *rec-8* mutants (Severson et al., 2009). To confirm that both types of cohesin promote axis integrity in pachytene nuclei, we used CRISPR to introduce the 3XTEV motifs in *rec-8* and *coh-3* in a strain carrying a *coh-4* null mutation. TEV injection in these germlines induced efficient removal of both REC-8 and COH-3 from axial elements (Figure 2D) and caused loss of HTP-3 tracks, confirming that both REC-8 and COH-3 cohesin promote axis stability in pachytene. We further tested this by depleting SMC-1, which should affect all types of cohesin, in pachytene nuclei using the auxin degron system (Zhang et al., 2015). This method induced slower and less penetrant cohesin depletion from axial elements compared with the TEV approach. After 8 hours of auxin treatment we observed partial SMC-1 removal from axial elements, but importantly regions lacking SMC-1 also lacked HIM-3 and SYP-1, confirming a local requirement for cohesin in axis and SC stability (Figures S2D-E). After 16 hours of auxin treatment we observed full disassembly of axial elements and the SC, with just a few short tracks remaining in late pachytene nuclei (Figure 2E). These results are consistent with REC-8 and COH-3/4 cohesin contributing independently to axis stability in pachytene nuclei.

### Cohesin removal reactivates chromosome movement and DSB formation in pachytene nuclei

CHK-2 acts as a master regulator of early prophase events in *C. elegans* (MacQueen and Villeneuve, 2001). Its activation at meiotic onset triggers DSB formation and chromosome movements that play a central role in homologue pairing and that induce chromosome clustering, giving chromatin a characteristic crescent shape appearance (Figures 1A-B). In wild-type germlines, chromosome clustering is restricted to nuclei in the transition zone, corresponding to leptotene/zygotene, as pachytene entry is marked by chromosome dispersal (Figure 1A). However, DAPI staining of germlines dissected 3.5 hours post REC-8 or COH-3 removal revealed nuclei with clustered chromosomes throughout most of the pachytene region (Figures 3A-B and Figure S3A), suggesting *de novo* establishment of chromosome movement. To confirm this, we used molecular markers of chromosome-end attachment to the nuclear envelope, a widely conserved feature of early prophase that in worms requires CHK-2 and pairing center-binding (PCB) proteins that localise to one end of each chromosome (Hillers et al., 2017). CHK-2 triggers recruitment of Polo-like kinase 2 (PLK-2) to PCB proteins by phosphorylating their Polo-binding domain, PLK-2 then promotes aggregation of SUN-1/ZYG-12 on the nuclear envelope to initiate cytoskeleton-driven chromosome movements (Harper et al., 2011; Labella et al., 2011; Penkner et al., 2009; Sato et al., 2009). In wild-type germlines, nuclei undergoing chromosome movements in transition zone display multiple PLK-2 aggregates, which disappear as nuclei progress into pachytene. Since a single PLK-2 aggregate associated with the paired X chromosomes often persists into pachytene until all chromosomes form CO precursors (Woglar et al., 2013), we used the presence of more than one PLK-2 aggregate as an indicator of active chromosome movement. In control germlines, nuclei with multiple PLK-2 aggregates correspond to approximately 30% of the total rows of nuclei progressing from leptotene to late pachytene (Figures 3A-B). In contrast, 3.5 hours post TEV-induced removal of REC-8^3XTEV^::GFP or COH-3^3XTEV^::mCherry, nuclei with multiple PLK-2 aggregates occupied over 70% of meiotic prophase, including most of the pachytene region (Figures 3A-B). We observed the same situation using antibodies against HIM-8 pT64, which label CHK-2-dependent phosphorylation of all PCB proteins at their Polo-binding domain (Kim et al., 2015), confirming re-acquisition of this CHK-2 marker in pachytene nuclei (Figure S3D-E). To determine if *de novo* formation of PLK-2 aggregates upon cohesin removal was functional, we monitored the presence of PLK-2-dependent SUN-1 S12 phosphorylation, which is normally restricted to transition zone nuclei (Harper et al., 2011; Labella et al., 2011). Similar to our observations above, removal of REC-8^3XTEV^::GFP or COH-3^3XTEV^::mCherry induced *de novo* appearance of SUN-1 pS12 in pachytene nuclei (Figures S3B-C). Therefore, removal of REC-8 or COH-3/4 from pachytene chromosomes induces rapid reacquisition of CHK-2-dependent markers of chromosome movement.

**Figure 3.**
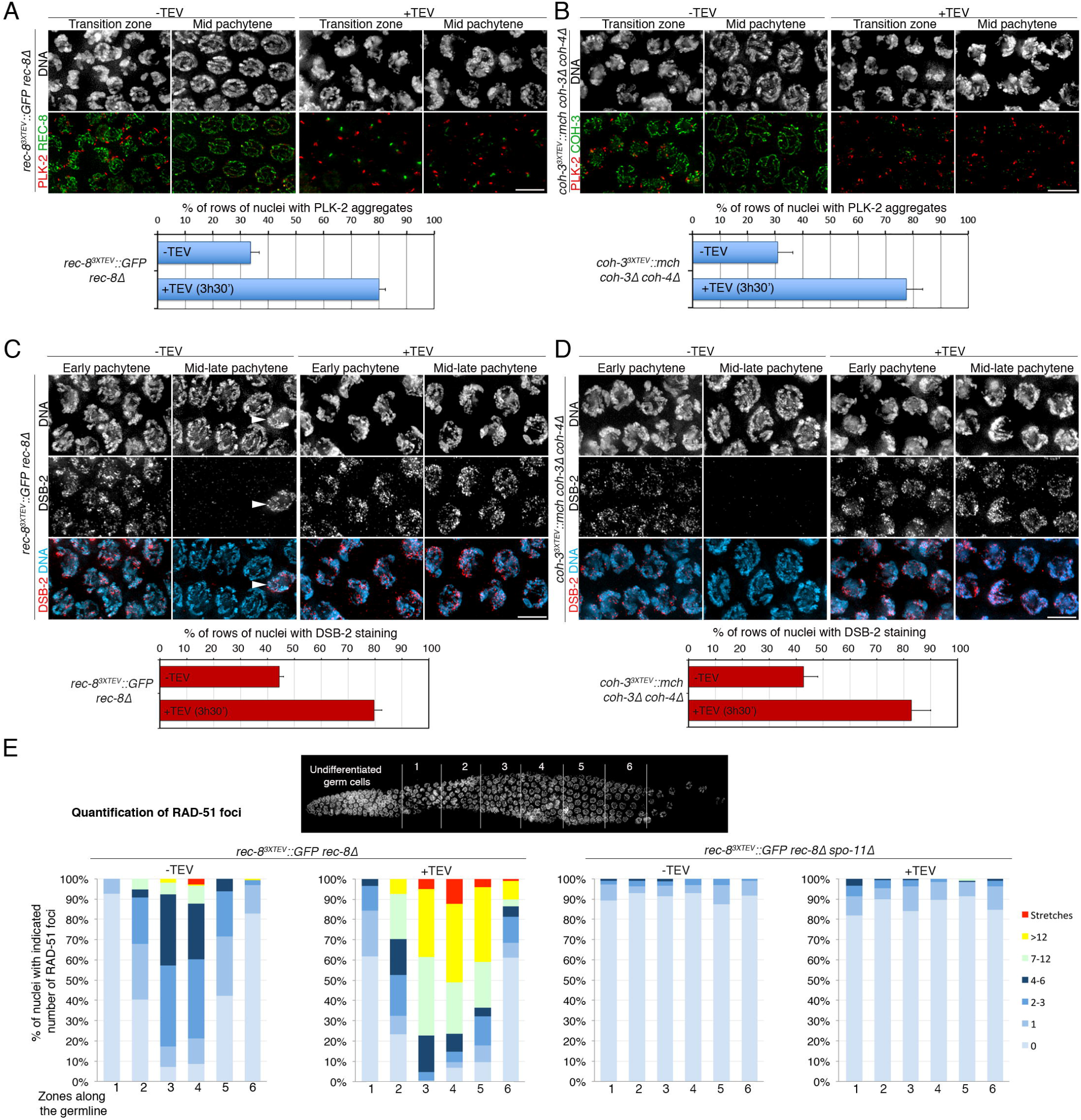
Cohesin removal induces reappearance of chromosome-movement and DSB-formation markers in pachytene nuclei. **(A-B)** TEV-mediated removal of REC-8^3XTEV^::GFP or COH-3^3XTEV^::mCherry induces reappearance of PLK-2 aggregates on the nuclear envelope. Quantification indicates percentage of rows of nuclei between meiotic onset and end of pachytene in which over 50% of nuclei contained more than 1 PLK-2 aggregate on the nuclear envelope. **(C-D)** TEV-mediated removal of REC-8^3XTEV^::GFP or COH-3^3XTEV^::mCherry induces reappearance of DSB-2 in pachytene nuclei. Arrow head on panel C indicates an arrested nucleus that retains DSB-2 in the pachytene region. Quantification indicates percentage of rows of nuclei between meiotic onset and end of pachytene in which over 50% of nuclei were positive for DSB-2. **(E)** Quantification of RAD-51 foci per nucleus before and after TEV-mediated removal of REC-8^3XTEV^::GFP in wild-type and *spo-11* mutant germlines (see table S1 for number of nuclei quantified per zone). The region between meiosis onset and the end of pachytene was divided into 6 zones as indicated in the DAPI-stained germline. Note that TEV injection increases RAD-51 foci in pachytene nuclei of wild-type, but not *spo-11* mutant, germlines. Scale bar= 5 μm in all panels. Three germlines were quantified for graphs in A-D and error bars represent standard deviation. See also Figure S3.

We next asked whether cohesin removal would also induce redeployment of the DSB machinery in pachytene nuclei. During early prophase, CHK-2 promotes association of DSB-1/2 with chromosomes, two factors required for DSB formation and whose association with chromosomes indicates a DSB-permissive state (Rosu et al., 2013; Stamper et al., 2013). In control germlines, DSB-2 staining was observed in transition zone and early-mid pachytene nuclei (Figures 3C-D). Removal of REC-8^3XTEV^::GFP or COH-3^3XTEV^::mCherry caused a clear increase in the percentage of rows of prophase nuclei positive for DSB-2 (Figures 3C-D), confirming *de novo* association of DSB-2 with pachytene chromosomes and suggesting reacquisition of DSB competence. Finally, we investigated the effect of REC-8 removal on RAD-51 foci, which label early recombination intermediates produced upon resection of SPO-11 DSBs (Colaiacovo et al., 2003). REC-8 removal induced increased RAD-51 foci at all prophase stages (up to late pachytene), and this increase was dependent on SPO-11, as RAD-51 foci were largely absent when REC-8 was removed in a *spo-11* mutant background (Figure 3E and Figure S3F). The increase in RAD-51 foci induced by REC-8 removal can be explained by a requirement of REC-8 cohesin in ongoing repair of DSBs generated before TEV injection and by the occurrence of *de novo* DSB formation, as suggested by the redeployment of DSB-2 to pachytene chromosomes.

These results demonstrate that pachytene nuclei with full SC and a normal complement of CO precursors retain the potential to reactivate early, CHK-2-dependent, prophase events and suggest that cohesin plays a central role in preventing untimely CHK-2 activation once nuclei transition to pachytene.

### Direct SC removal from pachytene nuclei induces rapid reactivation of chromosome movement and DSB formation

The observation that cohesin removal from pachytene chromosomes triggers SC disassembly and reactivation of CHK-2-dependent events, led us to ask whether direct SC removal from pachytene chromosomes would on its own cause CHK-2 reactivation. We used the auxin-inducible system (Zhang et al., 2015) to attempt time-controlled degradation of the SC central region component SYP-2 in pachytene nuclei as we expected that removal of one component would be sufficient to destabilise the SC. Strikingly, four hours after young adult worms were placed on auxin plates, SC tracks, visualised with anti-SYP-1 antibodies, had disappeared from all pachytene nuclei (Figure 4A). Moreover, DAPI staining revealed reacquisition of chromosome clustering throughout most of the pachytene region, suggesting CHK-2 reactivation. Staining with anti-PLK-2 antibodies confirmed this, as over 70% of nuclei between leptotene and late pachytene displayed multiple PLK-2 aggregates in germlines from auxin treated worms, compared to 39% in untreated controls (Figure 4B and Figure S4A). *In vivo* imaging in worms expressing mScarlet::SYP-3 and PLK-2::GFP showed that SYP-2 depletion for just two hours triggered SC disassembly and reappearance of PLK-2 aggregates in pachytene nuclei (Figure 4F and Movies S1-3). Moreover, these experiments demonstrated that *de novo* formed PLK-2::GFP aggregates displayed extensive movement on the nuclear envelope, including fusion and splitting events characteristic of transition zone nuclei (Movie S4). SC depletion also triggered *de novo* association of DSB-2 with pachytene chromosomes (Figure 4C), and increased RAD-51 foci (Figures 4D-E), consistent with *de novo* DSB formation and/or impaired ongoing repair of previously formed DSBs. Interestingly, COSA-1 foci in late pachytene nuclei remained intact following complete SC disassembly (Figure S4B), consistent with previous reports that used 1,6-hexanediol to dissolve the SC in dissected germlines (Rog et al., 2017). Therefore, SC removal is sufficient to induce CHK-2 reactivation in pachytene nuclei, suggesting that SC surveillance remains active once the structure is fully assembled.

**Figure 4.**
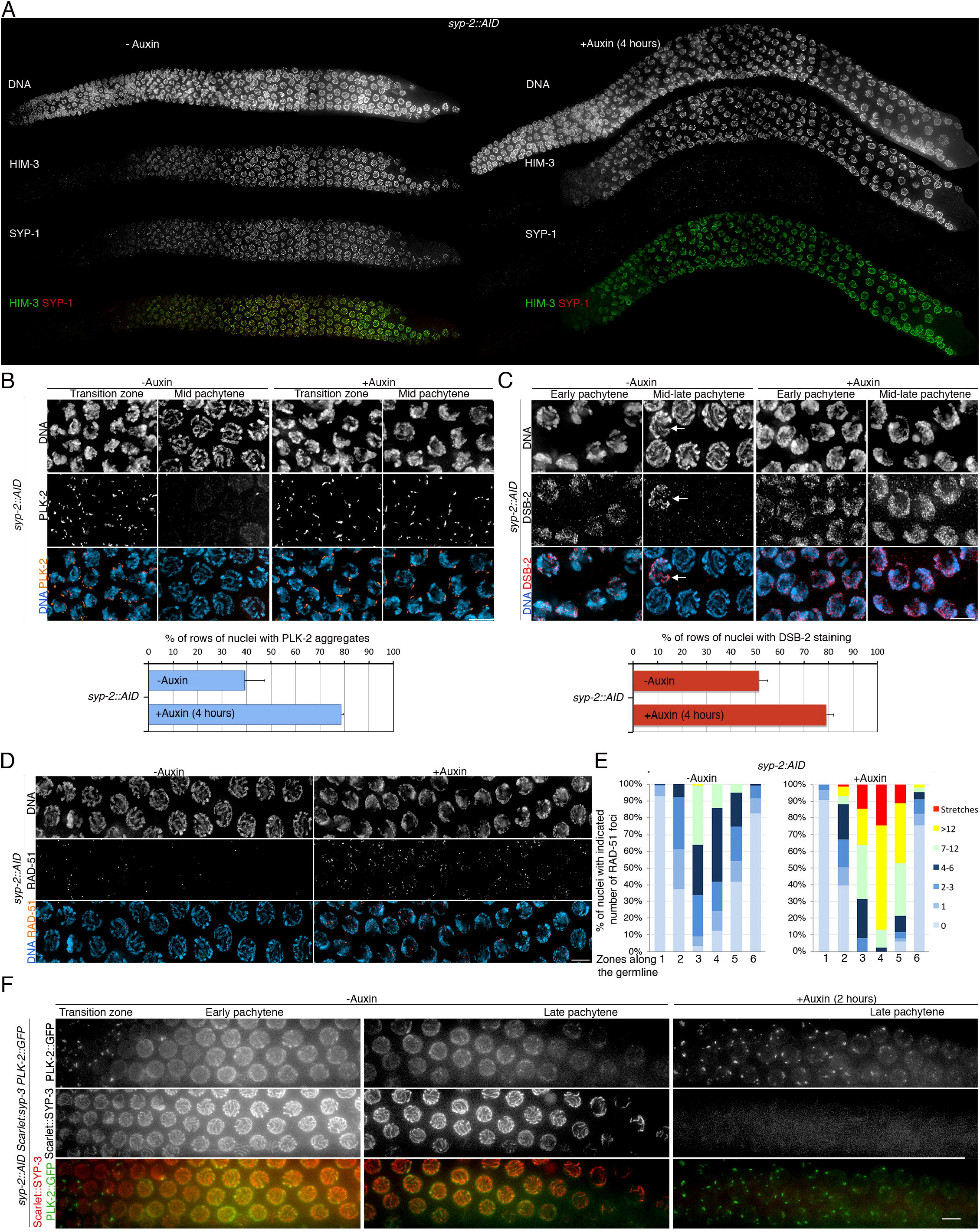
SC removal induces reappearance of chromosome-movement and DSB-formation markers in pachytene nuclei. **(A)** Auxin treatment of *syp-2::AID* worms induces rapid SC disassembly (visualised with anti-SYP-1 antibodies) without affecting axial elements (visualised with anti-HIM-3 antibodies). **(B-C)** Auxin treatment (4 hours) of *syp-2::AID* worms induces reappearance of PLK-2 aggregates on the nuclear envelope (B) and DSB-2 in pachytene nuclei (C). Quantification indicates percentage of rows of nuclei between meiotic onset and the end of pachytene in which over 50% of nuclei were positive for PLK-2 aggregates (more than 1 aggregate per nucleus) or DSB-2. **(D)** Auxin treatment (4 hours) of *syp-2::AID* worms induces accumulation of RAD-51 foci in pachytene nuclei. **(E)** Quantification of RAD-51 foci per nucleus before and after auxin-induced SC removal (see table S4 for number of nuclei quantified per zone). **(F)** *In vivo* imaging of germlines from *syp-2::AID* expressing mScarlet::SYP-3 (SC), PLK-2::GFP. Note SC disassembly and reappearance of PLK-2::GFP aggregates following 2 hours of auxin treatment. Scale bar= 5 μm in all panels. Three germlines were quantified for graphs in B-C and error bars represent standard deviation. See also Supplemental Movies 1-4 and Figure S4.

### Limited SC disassembly triggered by cohesin removal is sufficient for reactivation of CHK-2-dependent events

To further clarify the interplay between cohesin, SC stability, and CHK-2 reactivation we used different approaches to induce partial cohesin depletion from pachytene nuclei. First, we imaged *rec-8*^3XTEV^::*GFP* germlines that were dissected and fixed just 1.5 hours after TEV injection, instead of 3.5 hours as in previous experiments. Most mid pachytene nuclei from germlines dissected 1.5 hours post TEV injection displayed partial REC-8 removal and limited SC disassembly, but already displayed clear reappearance of PLK-2 aggregates associated with chromosomal ends (Figure S5A). Second, we monitored PLK-2 aggregates and the SC in germlines from *smc-1::AID::GFP* worms following 8 hours of auxin treatment, which induced partial cohesin removal (Figure S2E). These germlines also displayed partial SC disassembly in mid pachytene nuclei, which showed robust reappearance of PLK-2 aggregates (Figures S5B-D). Third, we also used the auxin degron system to deplete REC-8::AID::GFP from pachytene nuclei. Auxin treatment of these worms resulted in slower and less complete REC-8 removal compared with the TEV approach. 4 hours of auxin treatment induced partial loss of REC-8 and the SC from early pachytene nuclei, which displayed reappearance of PLK-2 aggregates (Figures S5E-F). By 8 hours of auxin treatment, PLK-2 aggregates also accumulated in mid pachytene nuclei (Figures S5F). These results suggest that the SC is locally destabilised when cohesin is removed from chromosomes and that this limited SC disassembly is sufficient to reactivate CHK-2 in a nucleus-wide fashion. Interestingly, in all three experimental approaches described above SC disassembly was much more pronounced in early pachytene nuclei than in mid and late pachytene nuclei, consistent with recent findings demonstrating the SC becomes more stable as nuclei progress through pachytene (Machovina et al., 2016; Nadarajan et al., 2017; Pattabiraman et al., 2017).

### Reinstallation of the DSB-formation and chromosome-movement machinery in pachytene nuclei requires CHK-2 and HORMA protein HTP-1

Our observations thus far show that cohesin and SC depletion from pachytene nuclei trigger reactivation of early steps of pairing and recombination, which under unchallenged conditions are restricted to early prophase stages and are dependent on CHK-2 activity. Thus, we sought to confirm whether the reinstatement of early prophase events seen in pachytene nuclei following cohesin removal occurs via CHK-2 activation. As *chk-2* mutants accumulate severe meiotic defects from the onset of meiosis, we tested whether the auxin system could be used to induce rapid CHK-2 depletion. Homozygous *chk-2::AID* worms (generated by CRISPR) displayed normal chiasma formation despite slower SC assembly. Importantly, following 6 hours of auxin treatment we observed complete loss of PLK-2 aggregates (Figure 5A) and DSB-2 staining (Figure 5B) from germlines *chk-2::AID* worms, consistent with loss of CHK-2 activity. Therefore, we treated *chk-2::AID* worms with auxin for 6 hours before inducing TEV-mediated REC-8^3XTEV^::GFP removal and evaluating the presence of PLK-2 aggregates on the nuclear envelope (Figure 5C). Injection of the TEV protease following CHK-2 depletion resulted in REC-8 removal (Figure 5C) and SC disassembly (Figure 5E) from pachytene chromosomes, but failed to induce reappearance of PLK-2 aggregates on the nuclear envelope (Figure 5C), as observed in control REC-8^3XTEV^::GFP germlines (Figure 5D). These results confirm that CHK-2 is required for reimplementing early prophase events in pachytene nuclei following REC-8 removal and SC disassembly.

**Figure 5.**
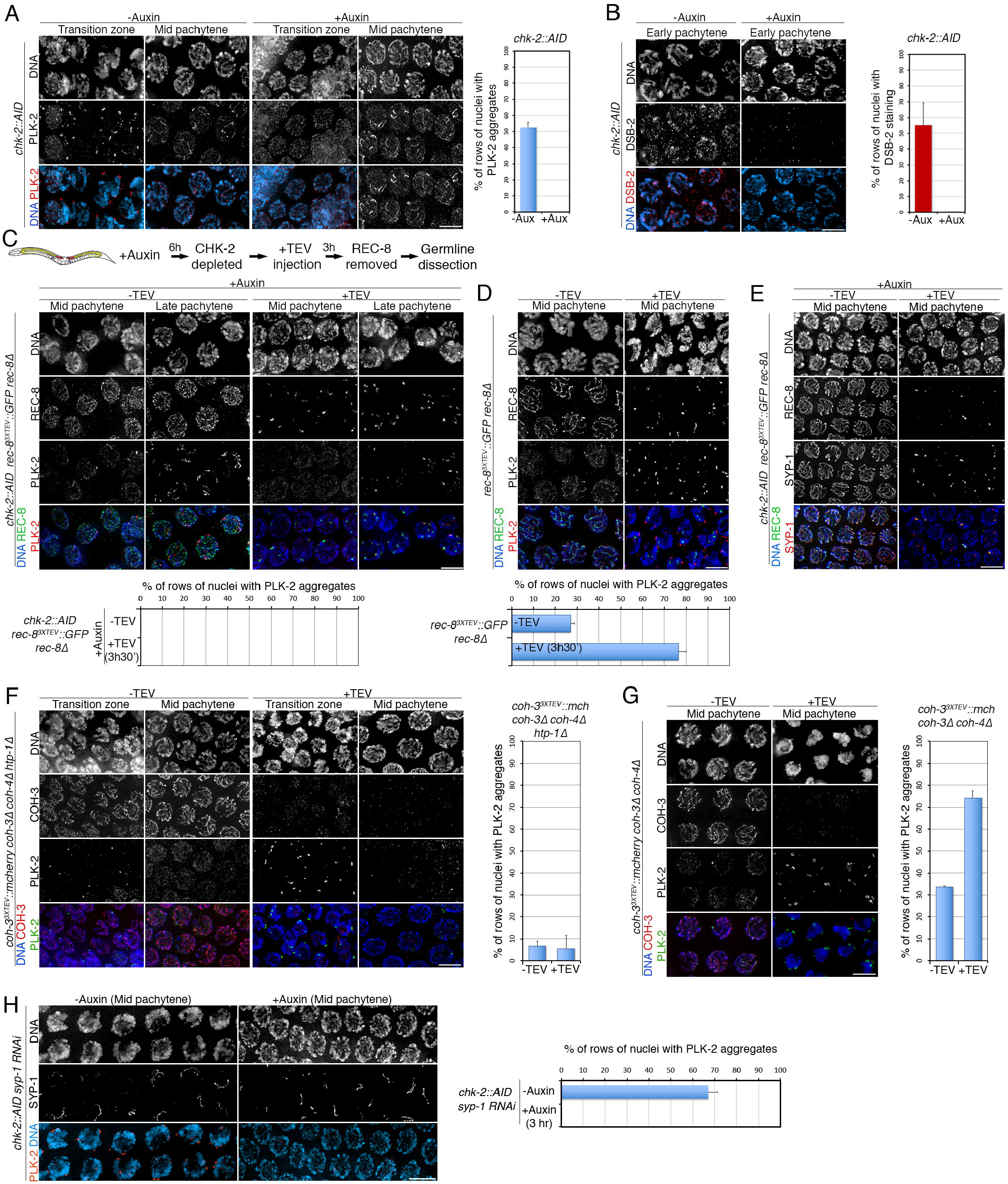
CHK-2 is required for reactivating chromosome movement and DSB formation in pachytene nuclei. **(A-B)** Auxin treatment (6 hours) of *chk-2::AID* worms induces disappearance of PLK-2 aggregates (A) and DSB-2 (B) from all stages of meiotic prophase. **(C-D)** Diagram of experimental design for auxin-induced CHK-2 depletion before TEV-mediated REC-8^3XTEV^::GFP removal. Note that in CHK-2-depleted germlines PLK-2 aggregates are not reformed following REC-8^3XTEV^::GFP removal (C). REC-8^3XTEV^::GFP removal induces reappearance of PLK-2 aggregates in pachytene nuclei (D). **(E)** TEV-mediated removal of REC-8^3XTEV^::GFP in CHK-2-depleted germlines induces SC disassembly. **(F-G)** TEV-mediated removal of COH-3^3XTEV^::mCherry in *htp-1* mutants fails to induce reappearance of PLK-2 aggregates in pachytene nuclei (F), as observed in wild-type germlines (G). **(H)** Auxin-mediated depletion of CHK-2::AID (3 hours) induces disappearance of PLK-2 aggregates from pachytene nuclei of *syp-1 RNAi* germlines. Graphs indicate percentage of rows of nuclei between meiotic onset and end of pachytene in which over 50% of nuclei contained more than 1 PLK-2 aggregate on the nuclear envelope (A and C-H), or DSB-2 staining (B). Scale bar= 5 μm in all panels. Three germlines were quantified for graphs in all panels and error bars represent standard deviation.

HORMA-domain protein HTP-1 is a key component of the feedback mechanisms that couple SC assembly with meiotic progression during early prophase (Couteau and Zetka, 2005; Martinez-Perez and Villeneuve, 2005). HTP-1 promotes persistence of CHK-2-dependent chromosome clustering in mutant backgrounds in which SC assembly fails in one or more chromosomes (Martinez-Perez and Villeneuve, 2005; Silva et al., 2014), presumably by participating in the creation and transmission of a signal that sustains CHK-2 activity in the presence of unsynapsed chromosomes. To investigate if HTP-1 is required for the reactivation of CHK-2-dependent events observed when cohesin is removed from pachytene chromosomes, we induced TEV-mediated removal of COH-3^3XTEV^::mCherry in germlines of *htp-1* mutants (carrying null alleles of endogenous *coh-3* and *coh-4*). TEV injection induced efficient COH-3^3XTEV^::mCherry removal in *htp-1* mutant germlines, but failed to induce reappearance of chromosome clustering and formation of multiple PLK-2 aggregates on the nuclear envelope of pachytene nuclei (Figure 5F), as observed when COH-3^3XTEV^::mCherry was removed in control germlines (Figure 5G). This suggests that the structural changes that result when cohesin is removed from pachytene chromosomes are sensed and transmitted by the same, HTP-1-dependent, surveillance mechanisms that monitor SC assembly to regulate CHK-2 activity during early prophase.

### CHK-2 depletion causes rapid disassembly of the DSB-formation and chromosome-movement machinery

During the control experiments to validate the depletion of CHK-2 using the auxin degron, we observed that auxin treatment for 6 hours induced complete loss of PLK-2 aggregates and DSB-2 staining from transition zone and early pachytene nuclei (Figures 5A-B). This suggested that CHK-2 is not simply required at the onset of prophase to kick-start pairing and recombination, but rather that sustained CHK-2 activity during early prophase is required to prevent premature disassembly of the pairing and recombination machinery. We further tested this by depleting CHK-2 from *syp-1* RNAi germlines, which accumulate nuclei with PLK-2 aggregates and chromosome clustering throughout most of the pachytene region due to impaired synapsis from the onset of meiosis. Indeed, CHK-2 depletion for only 3 hours triggered complete loss of PLK-2 aggregates and chromosome clustering from all prophase nuclei (Figure 5H), confirming that persistence of PLK-2 aggregates and chromosome clustering require sustained CHK-2 activity. These results suggest that activation and inactivation of CHK-2 rapidly regulates the localization of the pairing and recombination machinery, whose components remain responsive to CHK-2 status through most of pachytene.

### CHK-2 reactivation switches back early meiotic events in the absence of transcriptional changes

The drastic changes in nuclear organization observed when cohesin or the SC are removed from pachytene nuclei, which include reappearance of chromosome-associated markers of DSB formation and chromosome movement, led us to ask whether reestablishment of these events involved transcriptional changes of the corresponding genes. Thus, we performed whole-worm RNAseq to compare the transcriptome before and after 4 hours of auxin-mediated depletion of SYP-2::AID, which induced rapid accumulation of nuclei with CHK-2-dependent markers in the pachytene region (Figures 4B-C), focusing our analysis on a list of 335 meiotic genes (Hillers et al., 2017; Ortiz et al., 2014) (Table S2). We detected no significant changes in the expression of meiotic genes (including *dsb-2* and *chk-2*) following SYP-2 depletion, but observed increased expression of the proapoptotic factor *egl-1* (Figure 6A), suggesting that SYP-2 depletion triggers an apoptotic response, consistent with the increased levels of apoptosis observed in *syp-2* mutants (Colaiacovo et al., 2003). Therefore, the striking changes in nuclear organization caused by SC disassembly occur in the absence of obvious transcriptional changes, suggesting that CHK-2 substrates remain present in pachytene nuclei and that their localisation is modulated according to the status of CHK-2 activity. In agreement with this, we found that despite the large increase in chromosome associated DSB-2 signal triggered by SYP-2::AID depletion (Figure 4C), overall protein levels of DSB-2 remained unchanged before and after SYP-2 removal (Figure 6B).

**Figure 6.**
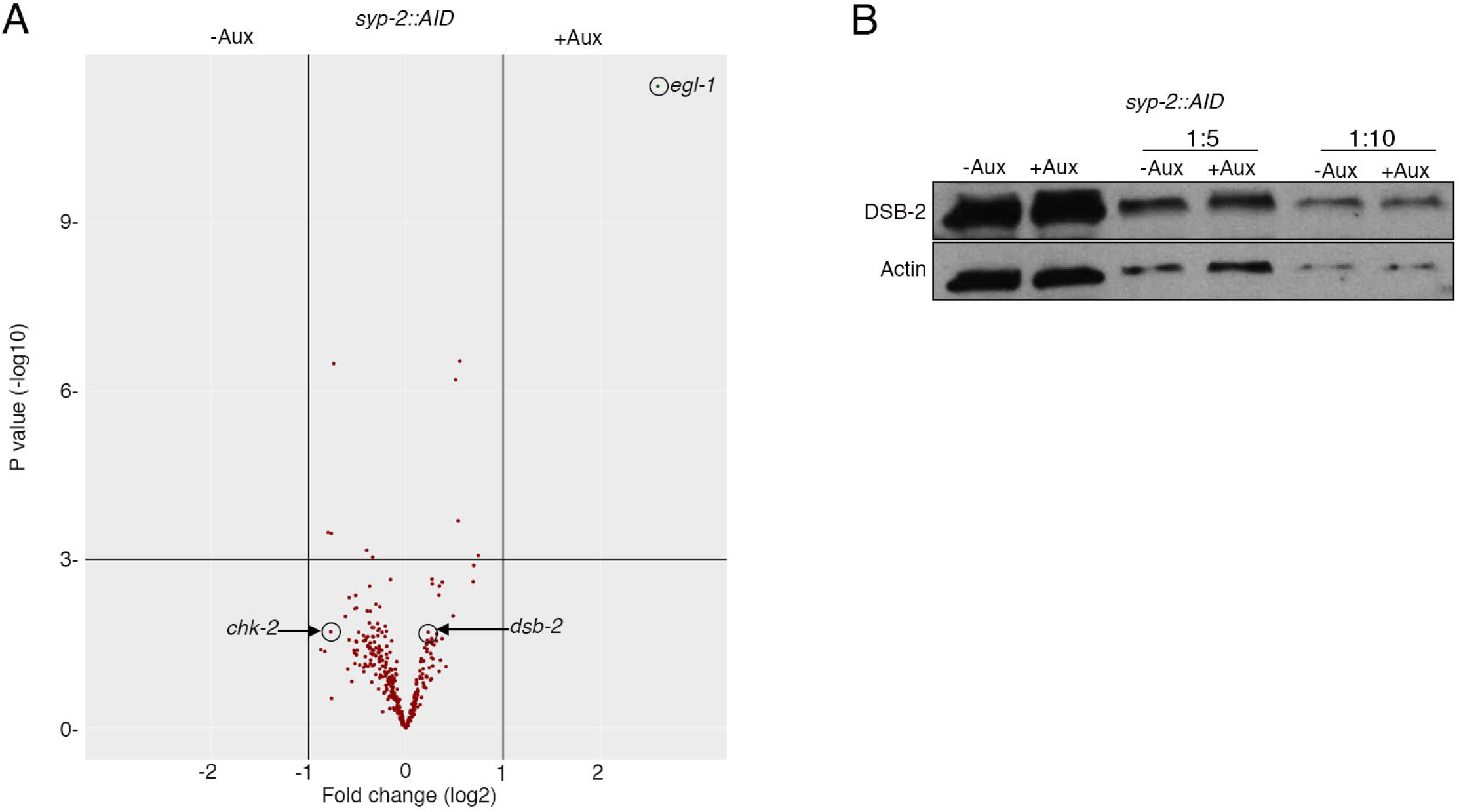
SC depletion does not trigger transcriptional changes of meiotic genes. **(A)** Volcano plot showing transcriptional changes (whole worm RNAseq) of 335 meiotic genes in *syp-2::AID* worms before and after (4 hours) auxin treatment. Note that *egl-1* is the only gene that appears significantly upregulated with a fold change higher than 2. **(B)** Western blot with extracts (1:1,1:5, and 1:10 dilution) from *syp-2:AID* worms before and after treatment showing that the amount of DSB-2 protein is not increased following SYP-2 depletion. Actin is used as loading control. See also Table S2.

### Cohesin-containing axial elements contribute to maintain CHK-2 activation

While performing experiments involving the simultaneous removal of COH-3^3XTEV^ and REC-8^3XTEV^ shown in Figure 2D, we noticed that reacquisition of chromosome clustering in pachytene nuclei 3.5 hours post TEV injection was less pronounced than when REC-8 or COH-3 were individually removed. This suggested that the integrity of cohesin-containing axial elements is important to fully reactivate CHK-2, or to sustain this activation. In fact, while we consistently observed the formation of multiple PLK-2 aggregates on the nuclear periphery of pachytene nuclei when REC-8 or COH-3 were individually removed (Figures 3A-B), or when SYP-2 was depleted (Figure 4B), simultaneous removal of REC-8^3XTEV^ and COH-3^3XTEV^ resulted in many early pachytene nuclei displaying a single PLK-2 aggregate associated with the nuclear periphery (Figure 7A). To clarify if cohesin-containing axial elements are required to sustain high levels of CHK-2 activity, we performed simultaneous removal of COH-3^TEV^ and REC-8^TEV^ in germlines of *syp-1* RNAi worms, in which impaired synapsis induces accumulation of nuclei with CHK-2 markers through most of the pachytene region. The efficiency of REC-8^3XTEV^ and COH-3^3XTEV^ removal was confirmed by the disappearance of HTP-3 tracks. As expected, control (*syp-1* RNAi) germlines displayed extensive accumulation of nuclei with multiple PLK-2 aggregates, while TEV injection caused a clear reduction in the number of these nuclei and the appearance of nuclei with 1 or no PLK-2 aggregates (Figure 7B and Figure S6). This confirms that the integrity of axial elements is required to sustain full CHK-2 activation. Interestingly though, since we observed complete loss of PLK-2 aggregates when we depleted CHK-2 from *syp-1* RNAi germlines (Figure 5H), the persistence of a single PLK-2 aggregate in many pachytene nuclei following the simultaneous removal of REC-8^3XTEV^ COH-3^3XTEV^ from *syp-1* RNAi germlines suggests that CHK-2 activity is not completely eliminated in this case, despite large axis disassembly.

**Figure 7.**
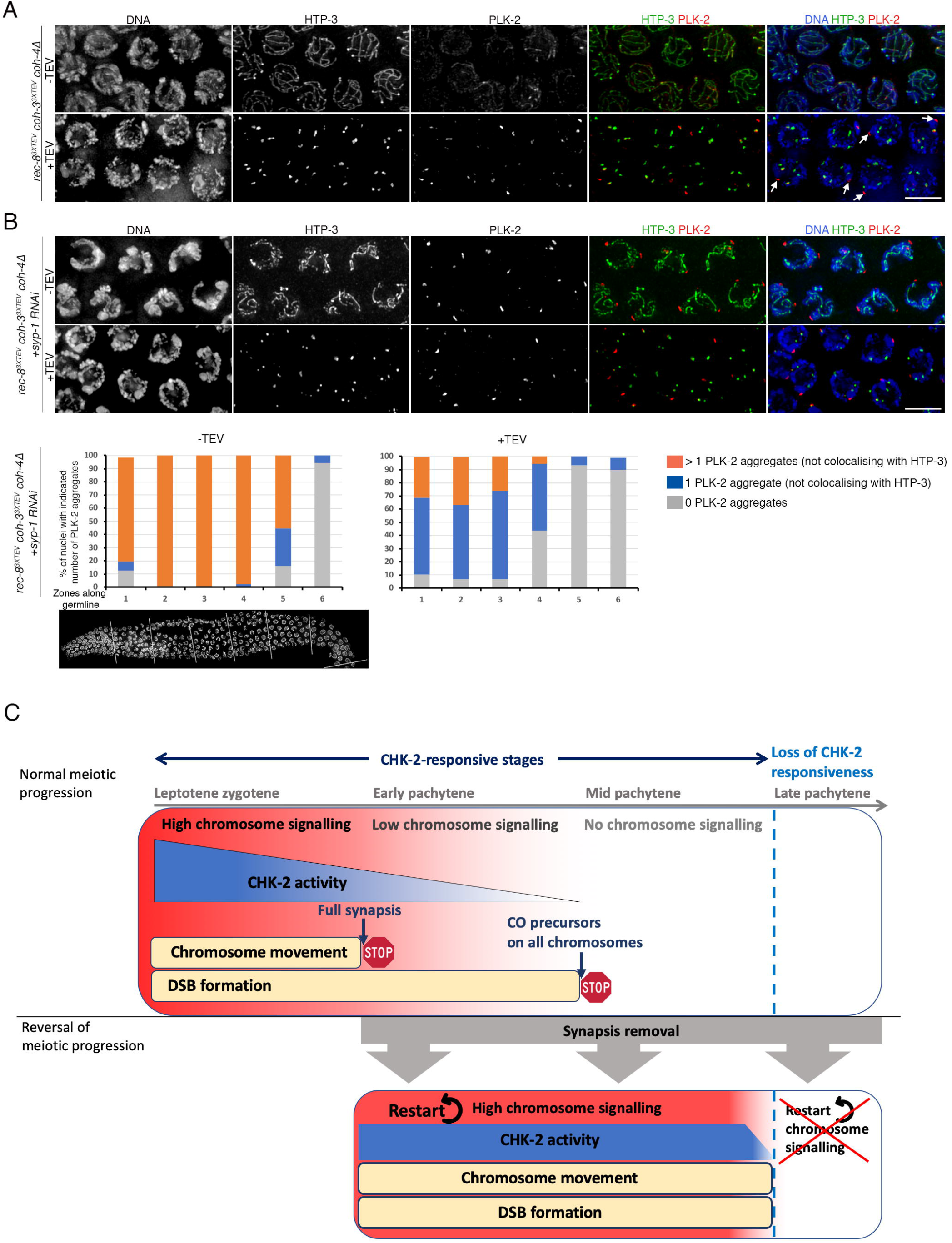
Axial elements contribute to sustain CHK-2 activity and model of CHK-2 feedback. **(A)** Simultaneous removal of REC-8^3XTEV^ and COH-3^3XTEV^ induces limited formation of PLK-2 aggregates on the nuclear envelope (not colocalising with nuclear aggregates of HTP-3). Compare to images in Figures 3A-B where either REC-8 or COH-3 are removed and multiple PLK-2 aggregates form on the nuclear envelope of pachytene nuclei **(B)** Simultaneous removal of REC-8^3XTEV^ and COH-3^3XTEV^ from *syp-1 RNAi* germlines reduces the number of PLK-2 aggregates on the nuclear envelope. Graphs show quantification of the percentage of nuclei with indicated number of PLK-2 aggregates, germlines were divided into six equal-size regions between meiotic onset and the end of pachytene and three germlines were scored for each condition. Scale bar= 5 μm in all panels. See also Figure S6 **(C)** Model: Synapsis-controlled chromosome signalling regulates meiotic progression. During normal meiotic progression chromosome signalling (red to white gradient) promotes CHK-2 activity (blue to white gradient). The presence of unsynapsed chromosomes during leptotene-zygotene induces high chromosome signalling, which sustains high CHK-2 activity, chromosome movement and DSB formation. Full synapsis causes a reduction of chromosome signalling and CHK-2 activity that terminates chromosome movement (first STOP sign). The formation of crossover precursors on all homologue pairs stabilises the SC, terminating chromosome signalling, and shutting down CHK-2 activity and DSB formation (second STOP sign). Removal of the SC from early and mid pachytene nuclei, either directly or by cohesin removal, restarts high chromosome signalling, inducing full CHK-2 activity and reactivating chromosome movement and DSB formation. Nuclei that transition to late pachytene lose the capacity to reactivate CHK-2.

### Late pachytene nuclei are unable to reactivate CHK-2 upon SC or cohesin removal

In all the experiments described above we noted that cohesin removal (REC-8^3XTEV^ or COH-3^3XTEV^ individually removed) or SC depletion consistently induced reappearance of CHK-2 markers through most of the pachytene region, except for very late pachytene nuclei. For example, we typically observed that 70-80% of nuclei between leptotene and late pachytene became positive for PLK-2 aggregates and DSB-2 staining following cohesin removal or SC depletion, with the remaining 20-30% of nuclei lacking these CHK-2-dependent markers invariably corresponding to the rows of nuclei situated in late pachytene (Figures 3A-D and 4B-C). In fact, simultaneous visualization of PLK-2 aggregates and COSA-1 foci following REC-8 removal or SC depletion revealed few nuclei displaying both markers (Figure S7). As SC depletion in late pachytene nuclei occurs as efficiently as in earlier stages (Figure 4A), the inability to reactivate CHK-2-dependent events during late pachytene suggests that nuclei at this stage lose the ability to respond to the presence of unsynapsed chromosomes.

## Discussion

Our findings reveal that nuclei with fully synapsed chromosomes and a normal complement of CO precursors undergo rapid, nucleus-wide, reestablishment of chromosome movement and DSB formation when SC stability is compromised. This process requires *de novo* CHK-2 activation and causes a dramatic reorganization of the nucleus that appears to functionally revert pachytene nuclei back to earlier prophase stages, revealing a striking plasticity of the meiotic program. We identify SC surveillance as the mechanism that regulates CHK-2 activity during pachytene. Moreover, we also identify a key role for REC-8 and COH-3/4 cohesin in promoting SC stability, and therefore meiotic progression, as removal of either type of cohesin causes rapid SC disassembly and CHK-2 reactivation in pachytene nuclei. Below, we discuss our findings in the context of a model in which SC surveillance orchestrates meiotic progression by continuously regulating CHK-2 activity (Figure 7C).

### Meiotic progression requires continued regulation of CHK-2 activity

In *C. elegans* CHK-2 promotes chromosome movement, DSB formation, and SC assembly at meiotic entrance (MacQueen and Villeneuve, 2001) and CHK-2 activity is lost gradually; markers of chromosome movement disappear at pachytene entrance, coinciding with full synapsis, while markers of DSB formation disappear at mid pachytene, presumably once all chromosomes have formed CO precursors (Rosu et al., 2013; Stamper et al., 2013; Woglar et al., 2013). In contrast, mutants with defects in synapsis or CO formation display persistent CHK-2 activity, evidencing feedback regulation of CHK-2 (Kim et al., 2015; Rosu et al., 2013; Stamper et al., 2013; Woglar et al., 2013). Synapsis deficient mutants arrest at leptotene/zygotene, retaining CHK-2-dependent markers of chromosome movement and DSB formation, and mutants deficient in CO formation but competent in SC assembly arrest at early pachytene, retaining only CHK-2-dependent markers of DSB formation (MacQueen et al., 2002; Rosu et al., 2013; Sato et al., 2009; Stamper et al., 2013). These and other observations suggest that the termination of chromosome movement at early pachytene and of DSB formation at mid pachytene, which are controlled by checkpoint mechanisms (Martinez-Perez and Villeneuve, 2005; Rosu et al., 2013; Stamper et al., 2013; Woglar et al., 2013), represent two key functional transitions of meiotic progression that culminate with a nucleus-wide loss in competency for CHK-2-dependent events. However, we now show that nuclei that have progressed normally to mid pachytene undergo rapid reactivation of CHK-2-dependent events, including chromosome movement, when SC stability is compromised, a finding that provides important insights into the surveillance mechanisms that regulate meiotic progression.

First, the fact that mid pachytene nuclei retain full potential for CHK-2 reactivation evidences that the kinase remains present in the nucleus in an inactive, but responsive, status and suggests that CHK-2 inactivation is actively sustained. In principle, CHK-2 inactivation could be achieved by inhibitory signals triggered when all chromosome pairs form CO precursors, or by the extinction of a positive signal that promotes CHK-2 activity until all chromosomes achieve synapsis and form CO precursors. Our findings support the latter hypothesis and suggest that the status of synapsis is key in extinguishing this positive signal (see below). Because late pachytene nuclei fail to reinstate markers of CHK-2 activity following forced SC removal, the transition to this stage must involve changes either in the capacity of chromosomes to generate the signal that feeds back to CHK-2 or in the ability of CHK-2 to respond to this signal. This observation is consistent with previous studies demonstrating that mutants that display meiotic arrest due to synapsis or recombination defects lose CHK-2 markers at late pachytene, despite persistence of the primary defect that induced the arrest (Kim et al., 2015; Martinez-Perez and Villeneuve, 2005; Woglar et al., 2013). We propose that meiotic prophase can be divided into two broad functional stages: one that spans between leptotene and late pachytene in which signalling from SC surveillance determines the nucleus-wide level of CHK-2 activity, which in turn controls chromosome movement and DSB formation, and a second that starts in late pachytene and is marked by a developmental transition that shuts down CHK-2 in a manner independent of SC surveillance (Figure 7C). The “CHK-2 responsive stage” can be further divided into three phases according to the level of CHK-2 activity present in the nucleus, which in wild-type germlines correlate with the sequential stages of early prophase. During leptotene/zygotene, high CHK-2 activity promotes chromosome movement, DSB formation, and SC assembly. Then, by early pachytene, once SC assembly is completed, CHK-2 activity decreases to levels that sustain DSB formation but not chromosome movement. Finally, by mid pachytene, once CO precursors are present on all chromosomes CHK-2 activity abates, but nuclei remain competent for CHK-2 reactivation. We propose that similar to Plk1 during mitosis in mammalian cells (Lera and Burkard, 2012), specific CHK-2-dependent functions may require different thresholds of kinase activity.

Second, our results uncover the presence of strong CHK-2-counteracting activities in prophase nuclei, which were evident by the rapid disappearance of markers of chromosome movement and DSB formation following CHK-2 depletion. This CHK-2 antagonistic activity is likely executed by phosphatases that dephosphorylate CHK-2 substrates to promote disassembly of the chromosome movement and DSB machineries. In addition, phosphatases could also antagonise CHK-2 directly by promoting dephosphorylation of activating sites, as seen in other organisms (Zannini et al., 2014). Identifying new CHK-2 substrates will be important to elucidate how the activity of CHK-2 and opposing phosphatases is integrated with the progression of pairing and recombination.

Third, *de novo* assembly of the chromosome-movement and DSB-formation machinery in the absence of transcriptional changes suggests that components of these complex processes remain present in the nucleus and are responsive to CHK-2 status until late pachytene. Consistent with this, we have shown that SC disassembly causes *de novo* phosphorylation of HIM-8 pT64, a CHK-2-dependent phosphoepitope that is present on the PC binding proteins of autosomes and X chromosome and promotes PLK-2 recruitment (Kim et al., 2015). Moreover, the presence of multiple HIM-8 pT64 (and PLK-2) aggregates following SC disassembly in mid pachytene nuclei reveals *de novo* loading of autosomal PCB proteins, as under normal progression these are unloaded at zygotene exit (Phillips and Dernburg, 2006). How specific substrates are affected by nucleus-wide levels of CHK-2 activity probably depends on complex interactions between CHK-2 and opposing phosphatases, which activity and subnuclear location are also likely to be highly regulated, as observed in the case of mitotic kinases and phosphatases (Gelens et al., 2018).

Fourth, our findings suggest that throughout the “CHK-2 responsive stage” the level of nucleus-wide CHK-2 activity is largely determined by the intensity of positive feedback signals emitted from chromosomes. HORMA proteins bound to axial elements, which can monitor *in situ* the status of pairing and recombination, are obvious candidates for signal generation (Kim et al., 2015), while a soluble pool of these proteins could also participate in the transmission of the signal to generate a nucleus-wide response (Silva et al., 2014). Our findings support the key role of HORMA proteins in signal generation/transmission as HORMA protein HTP-1 is required for reactivating CHK-2-dependent chromosome movement in pachytene nuclei following cohesin removal. How CHK-2 is activated in *C. elegans* remains unclear, as it lacks the N-terminal SQ/TQ cluster that in other organisms is phosphorylated by ATM and ATR to induce CHK-2 activation (Zannini et al., 2014), but it may involve dimerization and autophosphorylation in trans in its activation loop, as observed in yeast Mek1 (Niu et al., 2007). CHK-2 deactivation could be mediated by phosphorylation removal at activating sites, or by phosphorylation of inactivating sites at its FHA domain, as observed in mammalian Chk2 (van Vugt et al., 2010). Elucidating how signals from chromosomes and the activity of phosphatases are integrated to implement nucleus-wide control of CHK-2 remains an important goal for future studies.

### SC surveillance as a nucleus-wide regulator of CHK-2 activity

Two key aspects of our model for feedback regulation of CHK-2 are that surveillance of the chromosomal features responsible for signal generation must operate continuously until late pachytene, and that signal emission can be reactivated if SC stability is compromised during pachytene. *C. elegans* mutants with defects in synapsis arrest at earlier stages (leptotene/zygotene) than mutants deficient in CO formation but competent in SC assembly, which arrest at early pachytene, suggesting that synapsis and CO formation may be differently monitored to feedback on CHK-2 (Stamper et al., 2013; Woglar et al., 2013). However, recent studies show that the state of the SC is modified by the formation of CO precursors (Machovina et al., 2016; Nadarajan et al., 2017; Pattabiraman et al., 2017), opening the possibility that SC state itself rather than a specific recombination intermediate may be the feature under surveillance to monitor formation of CO precursors (Pattabiraman et al., 2017). By showing that the formation of CO precursors induces PLK-2-dependent phosphorylation of SC component SYP-4, which spreads along the whole length of the SC increasing its stability, (Nadarajan et al., 2017) provided a mechanistic link between recombination and SC status. As this mechanism operates in a chromosome autonomous manner, surveillance of SC status can explain how a single homologue pair lacking a CO precursor will keep emitting signals that sustain CHK-2 activity and how such signal will be extinguished once all homologue pairs form CO precursors. Our findings support a model in which meiotic progression is controlled by surveillance of synapsis status. We propose that during early prophase, or when SC stability is compromised in pachytene nuclei, unsynapsed axial elements emit signals that induce/sustain high CHK-2 activity, while synapsed axial elements in the context of homologues lacking a CO precursor emit signals that result in lower CHK-2 activity, sustaining DSB formation but not chromosome movement. Then, recombination-dependent SC stabilization terminates, albeit in a reversible manner, axis signalling. Our studies suggest that the small number of nuclei displaying chromosome clustering and CHK-2 markers of early prophase in the pachytene region of wild type germlines (arrowhead in Fig 3C), which are thought to be nuclei arrested at early prophase, may represent nuclei in which unscheduled SC disassembly/destabilisation, for example during interlock resolution, triggered CHK-2 reactivation. Our model relies on continued SC monitoring between leptotene and late pachytene, which is possible in *C. elegans* because HORMA proteins remain bound to axial elements throughout pachytene. Mammals also use HORMAD proteins to monitor synapsis (Wojtasz et al., 2012), but in contrast to *C. elegans* synapsis triggers ejection of HORMAs from axial elements (Wojtasz et al., 2009), thus terminating axis signalling. Despite these differences in HORMA proteins behaviour, the role of the SC in coupling the formation of CO precursors to the termination of axis signalling may be conserved between worms and mammals, as SC assembly is recombination dependent in mammals.

Studies in different organisms show that synapsis contributes to implement local and chromosome-wide regulation of events such as DSB formation, DSB repair, and CO formation (Kauppi et al., 2013; Libuda et al., 2013; Subramanian et al., 2016). Our study highlights how SC surveillance contributes to integrate pairing and recombination with meiotic progression by ensuring timely and sustained termination of chromosome movement and DSB formation in a nucleus-wide fashion. This role of the SC must be coordinated with its roles in promoting CO formation locally at CO-designated sites (Woglar and Villeneuve, 2018), and in limiting DSB and CO formation along paired homologues (Libuda et al., 2013; Nadarajan et al., 2017). Thus, signalling coupled to the SC operates in a chromosome-autonomous and nucleus-wide manner to regulate DSB and CO formation, control nuclear organization, and promote meiotic progression.

## Supporting information

Detailed methods are provided in the Supplemental Information

## Acknowledgements

We thank Laurence Game and Ivan Andrew from the MRC LMS Genomics facility for library preparation for RNAseq experiments, the *Caenorhabditis* Genetics Center (CGC) for providing *C. elegans* strains, and the following individuals for providing antibodies: Anne Villeneuve, Monique Zetka, Verena Jantsch, and Rueyling Lin. This work was supported by an MRC core-funded grant to E.M.-P. and by postdoctoral Fellowships from Fundación Alfonso Martin Escudero and EMBO to M.C.-P.

## Methods

Detailed methods are provided in the Supplemental Information.

