## Supplementary material for "Reactivation of chromosome signalling induces reversal of the meiotic program": Detailed methods are provided in the Supplemental Information

**Supplemental Figure Legends**

**Figure S1. (A)** Quantification of embryonic lethality in strains of indicated genotypes. Note that the *rec-8*$^{3XTEV}$*::GFP* and *coh-3*$^{3XTEV}$*::mCherry* transgenes rescue the high embryonic lethality of the *rec-8* single and *coh-3 coh-4* double mutants respectively. **(B)** Examples of diakinesis oocytes showing that expression of *rec-8*$^{3XTEV}$*::GFP* and *coh-3*$^{3XTEV}$*::mCherry* transgenes rescues chiasma formation in *rec-8* single and *coh-3 coh-4* double mutants respectively. **(C)** Injection of the TEV protease in germlines of worms expressing *rec-8::GFP* or *coh-3::mCherry* transgenes lacking the TEV recognition motif demonstrates normal cohesin localization and no apparent meiotic defects. The efficiency of the injection is confirmed by the incorporation of labelled nucleotides (Rho-dUTPs), which were co-injected with the TEV protease. Nuclei inside white rectangle are magnified on the inset shown below the germlines. **(D)** Whole germlines of indicated genotype before and after (3.5 hours) TEV protease injection demonstrating efficient removal of REC-8$^{3XTEV}$::GFP, while COH-3/4 remain associated with chromosomes. **(E)** Whole germlines of indicated genotype before and after (3.5 hours) TEV protease injection demonstrating efficient removal of COH-3$^{3XTEV}$::mCherry, while REC-8 remains associated with chromosomes. Scale bar= 5 µm in all panels.

**Figure S2. (A)** Whole germlines of indicated genotype before and after (3.5 hours) TEV protease injection showing that COH-3$^{3XTEV}$::mCherry removal induces SC disassembly. Note that some small SYP-1 signals persist colocalising with COH-3$^{3XTEV}$::mCherry signals that are more prominent in late pachytene nuclei, where their number is close to 6. Nuclei inside white rectangle are magnified on the inset shown below the germlines. **(B-C)** HORMA proteins HIM-3 (B) and HTP-1/2 (C) remain associated with axial elements following TEV-mediated removal of REC-8$^{3XTEV}$::GFP or COH-3$^{3XTEV}$::mCherry. **(D-E)** Auxin-mediated depletion of SMC-1::AID::GFP for 8 hours induces partial loss of cohesin from chromosomes. Note that regions lacking SMC-1 staining also lack HIM-3 and SYP-1 staining (arrowheads in E). Images were acquired using a structural illumination microscope. Scale bar= 5 µm in all panels.

**Figure S3. (A)** Whole germlines of indicated genotypes showing PLK-2 localization. Double-headed arrows indicates the beginning and end of regions containing nuclei with PLK-2 aggregates on the nuclear envelope. Note that in WT germlines PLK-2 aggregates are only present in the transition zone, while germlines stained 3.5 hours after TEV-mediated removal of REC-8$^{3XTEV}$::GFP or COH-3$^{3XTEV}$::mCherry contain nuclei with PLK-2 aggregates through most of the pachytene region. **(B-E)** TEV-mediated removal of REC-8$^{3XTEV}$::GFP or COH-3$^{3XTEV}$::mCherry induces reappearance of SUN-1 pS12 and HIM-8 pT64 signals on the nuclear envelope of pachytene nuclei. Graphs show percentage of rows of nuclei between meiotic onset and end of pachytene in which over 50% of nuclei contained more than 1 aggregate of SUN-1 pS12 or HIM-8 pT64 aggregate on the nuclear envelope. **(F)** TEV-mediated removal of REC-8$^{3XTEV}$::GFP induces accumulation of RAD-51 foci in pachytene nuclei. Three germlines were quantified per condition in graphs shown in (B-E) and error bars represent standard deviation. Scale bar= 5 µm in all panels.

**Figure S4. (A)** Whole germlines of *syp-2::AID* worms before and after (4 hours) auxin treatment, which induces appearance of PLK-2 aggregates on the nuclear envelope in most pachytene nuclei. **(B)** Late pachytene nuclei from *syp-2::AID; cosa-1::HA* worms showing that COSA-1 foci persist after 4 hours of auxin treatment. Graph shows quantification of COSA-1::HA foci in late pachytene nuclei from 5 germlines of *syp-2::AID; cosa-1::HA* worms before and after auxin treatment. Note that most nuclei show 6 foci before and after auxin treatment. Scale bar= 5 µm in all panels.

**Figure S5. (A)** Whole germline from *rec-8$^{3XTEV}$::GFP rec-8Δ* dissected 1.5 hours post TEV injection. Note partial REC-8 removal and SC disassembly (SYP-1) in mid pachytene nuclei that display multiple PLK-2 aggregates on the nuclear envelope. Nuclei inside white rectangle are magnified on the inset shown on the right-hand side of the panel. **(B-D)** 8 hours of auxin-mediated depletion of SMC-1::AID::GFP causes partial SC disassembly and appearance of PLK-2 aggregates in mid pachytene nuclei. Note that SC disassembly is more prominent in transition zone and early pachytene nuclei. Scale bar= 5 µm in all panels. Graph shows percentage of rows of nuclei between meiotic onset and end of pachytene in which over 50% of nuclei contained more than 1 PLK-2 aggregate. Three germlines were quantified per condition and error bars represent standard deviation. **(E)** Whole germline from *rec-8::AID::GFP rec-8Δ* dissected after 4 hours of auxin treatment. Note partial REC-8::AID::GFP depletion and SC disassembly in mid pachytene nuclei that display PLK-2 aggregates on the nuclear envelope. Nuclei inside white rectangle are magnified on the inset shown on the right-hand side of the panel. **(F)** Graph shows percentage of rows of nuclei between meiotic onset and end of pachytene in which over 50% of nuclei contained more than 1 PLK-2 aggregate after 0, 4, and 8 hours of auxin treatment. Three germlines were quantified per condition and error bars represent standard deviation. Scale bar= 5 µm in all panels.

**Figure S6.** Germlines from *rec-8$^{3XTEV}$ coh-3$^{3XTEV}$ coh-4Δ; syp-1RNAi* worms before (top) and after (bottom) TEV injection. Note intact HTP-3 axial elements and the presence of multiple PLK-2 aggregates on the nuclear envelope before TEV injection. TEV injection induces large disassembly of axial elements, visualised by the accumulation of HTP-3 in small nuclear aggregates, and a reduction both in the number of nuclei displaying PLK-2 aggregates as well as in the number of PLK-2 aggregates per nucleus.

**Figure S7. (A)** Whole germline from *rec-8$^{3XTEV}$::GFP rec-8Δ; cosa-1::HA* worms dissected 3.5 hours post TEV injection. Note that PLK-2 aggregates are lacking in late pachytene nuclei displaying bright COSA-1 foci. Nuclei inside white rectangle are magnified on the inset shown on the right-hand side of the panel. **(B)** Whole germline from *syp-2::AID; cosa-1::HA* worms dissected after 4 hours of auxin treatment. Note that PLK-2 aggregates are lacking in late pachytene nuclei displaying bright COSA-1 foci. Nuclei inside white rectangle are magnified on the inset shown on the right-hand side of the panel. Scale bar= 5 µm in all panels.

**Movie S1.** *In vivo* imaging of the transition zone and early pachytene regions of the germline from a *mScarlet::syp-3; syp-2::AID; fqSi13[PLK-2::GFP]* worm (not treated with auxin). Note the presence of dynamic PLK-2 aggregates (green) on the nuclear envelope of transition zone nuclei at the bottom left corner of the video and the presence of SC tracks (red) on all pachytene nuclei. Filming corresponds to 7 minutes.

**Movie S2.** *In vivo* imaging of the mid and late pachytene regions (left to right on the video) of the germline from a *mScarlet::syp-3; syp-2::AID; fqSi13[PLK-2::GFP]* worm (not treated with auxin). Note absence of PLK-2 aggregates (green) and the presence of SC tracks (red) on all nuclei. Filming corresponds to 7 minutes.

**Movie S3.** *In vivo* imaging of the mid and late pachytene regions (left to right on the video) of the germline from a *mScarlet::syp-3; syp-2::AID; fqSi13[PLK-2::GFP]* worm after 2 hours of auxin treatment. Note presence of dynamic PLK-2 aggregates (green) on the the nuclear envelop of most nuclei, apart from late pachytene nuclei at the bottom right-hand corner, and the absence of SC tracks (red) on all nuclei. Filming corresponds to 7 minutes.

**Movie S4.** Zoomed pachytene nucleus from video S3 (*mScarlet::syp-3; syp-2::AID; fqSi13[PLK-2::GFP]* after 2 hours of auxin treatment). Note that PLK-2 aggregates (green) undergo fusion and splitting events.

#### Methods

##### *C. elegans* strains and culture conditions

All strains were grown on *E. coli* (OP50) seeded NG agar plates at 20 °C under standard conditions. Unless otherwise indicated, all experiments were performed using young adults at 18-24 hours post L4. The N2 Bristol strain was used as wild-type strain. The following alleles were used: LG IV: *htp-1 (gk174), rec-8 (ok978), spo-11 (ok79);* LG V: *coh-3 (gk112), coh-4 (tm1857)*. Table S3 contains a full list of the strains generated in this study. Strains VC666 [*rec-8 (ok978) / nT1 [unc-? (n754) let-? qls50] (IV;V)*] and TY5120 [*coh-3(gk112) coh-4(tm1857) V / nT1 [unc-? (n754) let-? qls50] (IV;V)*] were used for viability assays.

##### Generation of transgenic *C. elegans* strains

Transgenic strains were generated using CRISPR to modify the endogenous locus or single-copy insertions of the desired transgene in strains carrying the MosCI transposon at the *ttTi5605* (chromosome II) or the *oxTi177* (chromosome IV) loci (Frokjaer-Jensen et al., 2008). The *rec-8$^{3XTEV}$::GFP* transgene was generated by adding a 75 bp fragment encoding for 3 repeats of the TEV recognition motif (ENLYFQGASENLYFQGELENLYFQG) after REC-8's Q289 codon in a vector expressing REC-8::GFP under the *rec-8* promoter and 3' UTR (Crawley et al., 2016). The *coh-3$^{3XTEV}$::mCherry* transgene was generated by adding a 75 bp fragment encoding for 3 repeats of the TEV recognition motif (ENLYFQGASENLYFQGELENLYFQG) after COH-3's I315 codon in a vector expressing COH-3::GFP under the *coh-3* promoter and 3' UTR. The *rec-8::AID::GFP* transgene was generated by adding a 135 bp fragment encoding the 35 amino acids of the AID tag (Zhang et al., 2015) before the start codon of GFP. The *plk-2::GFP* transgene was generated by fusing the endogenous sequence of *plk-2*, including introns, 860 bp of upstream and 1393 bp of downstream sequence to a GFP sequence containing three introns. Table S4 contains a list of all transgenes used in this study.

CRISPR-mediated genome editing was performed using preassembled Cas9-sgRNA complexes, single-stranded DNA oligos as repair templates, and *dpy-10* as a co-injection marker as described in (Paix et al., 2017). To generate TEV-cleavable versions of REC-8 and COH-3, a 75 bp fragment encoding for 3 repeats of the TEV recognition motif (ENLYFQGASENLYFQGELENLYFQG) was inserted after REC-8's Q289 and COH-3's I315 codons using a single-stranded DNA oligo as repair template (Table S5). To generate *chk-*

*2::AID* and *syp-2::AID* alleles we introduced 135 bp encoding the 35 amino acids of the AID tag (Zhang et al., 2015) before the stop codon of the genes using two single-stranded DNA oligos with a 35 bp overlap as repair templates (Table S4). We also used this strategy to generate an *smc-1::AID::GFP* allele by introducing the 135 bp of the AID tag before the start codon of GFP in the *smc-1* (*fq20[smc-1::GFP]*) allele that we generated previously (Crawley et al., 2016). Table S5 offers a full list of CRISPR alleles generated in this study.

**Auxin-mediated protein degradation**

All strains used for auxin-mediated protein degradation were homozygous for the *ieSi38* transgene expressing the TIR1 protein under the *sun-1* promoter, which confers efficient germline expression (Zhang et al., 2015). Auxin treatment was performed by placing young adult worms in seeded NG agar plates containing 1 or 4 mM Auxin for the indicated periods of time.

**TEV protease microinjection**

Germline injections were performed in young adult worms at 18-24 hours post L4 immobilized in 2% agarose pads and covered with Halocarbon oil 700 (Sigma) using a Narishige IM-31 pneumatic microinjector attached to an inverted Olympus IX71 microscope. Needles were made using borosilicate glass filaments with a 1.0 mm O.D. and 0.58 mm I.D. (BF100-58-10, Sutter Instruments) and a micropipette puller P-97 (Intracell). In all experiments except those displayed in Figure S2A, AcTEV™ Protease (Thermo Fisher, Cat. No. 12575) was used in a mix containing 10U/µl TEV protease in 50 mM Tris-HCl, pH 7.5, 1 mM EDTA, 5 mM DTT, 50% (v/v) glycerol, 0.1% (w/v) Triton X-100. For Figure S2A we used TEV protease from GenScript (Cat. No. Z03030-1000) in a final mix containing 2.5 ng/µl TEV protease in 50mM Tris, 5mM DTT, 12.5% glycerol, pH 7.5. In indicated experiments 25 pmol of tetramethyl-Rhodamine-5-dUTP (Roche) were added to the injection mix to evaluate the efficiency of germline microinjection by the incorporation of labelled nucleotides into the DNA of germ cells.  Following microinjection, worms were rescued from the agarose pad with M9 salt buffer and placed in NG plates with *E. coli* OP50 for 3h30' (unless otherwise indicated) before germline dissection.

**Immunostaining and image acquisition**

Germlines from 18-24 hours post L4 worms were dissected in EGG buffer (118 mM NaCl, 48 mM $KCl_2$, 2mM $CaCl_2$, 2mM $MgCl_2$, 5mM HEPES) containing 0.1% Tween and then fixed in the same buffer containing 1% paraformaldehyde for 5 minutes. Slides were immersed in liquid nitrogen before removing the coverslip and then placed in methanol at -20 $^{o}$C for 5 minutes. Slides were then washed three times for minutes each in PBST (1x PBS, 0.1% Tween) and then blocked in PBST containing 0.5% BSA for 1 hour. Slides were then incubated with primary antibodies diluted in PBST over night at room temperature. Following three washes of 10 minutes each in PBST, slides were incubated in the dark at room temperature for 2 hours with secondary antibodies diluted in PBST. All secondary antibodies were conjugated to Alexa-488, Alexa-555 or Alexa-647 (Life Technologies) and used at 1:500. Following three washes with PBST, slides were counterstained with DAPI, washed for 1h in PBST and mounted using Vectashield (Vector). Unless otherwise indicated, all images were acquired as three-dimensional stacks on a Delta Vision system equipped with an Olympus 1x70 microscope using a 100X lens. Images were subjected to deconvolution analysis using SoftWoRx 3.0 (Applied Precision) and images were mounted in Photoshop.

**Super resolution structured illumination microscopy**

Immunostaining was performed as described before, but slides were mounted using ProLong Diamond mounting media instead of Vectashield and covered using Zeiss High-performance 0.17 ± 0.005 cover slips. Images were acquired using a Zeiss Elyra S1 SIM microscope and mounted in Photoshop.

**Antibodies used in this study**

The following primary antibodies were used at the indicated dilutions: rabbit anti-GFP-488-conjugated (1:200) (Invitrogen), goat anti-GFP-488-conjugated (1:200) (Roche), rat anti-mCherry (1:1000) (5F8, Chromotek), rabbit anti-COH-3/4 (1:400) (Crawley et al., 2016), mouse anti-REC-8 (1:100) (Novus Biologicals), guinea pig anti-SYP-1 (1:400) (MacQueen et al., 2002), chicken anti-SYP-1 (1:300) (Silva et al., 2014), guinea pig anti-HTP-3 (1:800) (Goodyer et al., 2008), rabbit anti-HIM-3 (1:400) (Zetka et al., 1999), rabbit anti-HTP-1/2 (1:400) (Silva et al., 2014), rabbit anti-PLK-2 (1:500) (Nishi et al., 2008), SUN-1 pS12 (1:1000) (Woglar et al., 2013), rabbit anti-RAD-51 (1:10000) (Novus Biologicals), mouse anti-HA

(1:200) (Cell Signalling), rabbit anti-DSB-2 (1:1000) (Rosu et al., 2013). Phospho-specific antibodies against HIM-8 pT64 (1:400) were produced by injecting rabbits with the synthetic peptide DTPRFSpTPIVPNVC (GenScript). Polyclonal anti-HIM-8 pT64 antibodies were affinity purified by binding to a column containing the phospho-peptide and the specificity of the antibodies was validated by absence of staining in germlines of *chk-2* mutant worms.

**Quantification of RAD-51 foci**

Analysis of RAD-51 foci was performed as in (Silva et al., 2014).Each germline was divided into 6 equal-size regions, from the beginning of the transition zone (leptotene-zygotene) to the bend of the germline (end of pachytene, beginning of diplotene). The number of foci per nucleus in each region of the germline were quantified. When RAD-51 signals appeared as agglomerates instead of single foci they were categorized as stretches. At least three germlines per genotype and condition were used for quantification of RAD-51 foci and the number of nuclei counted for each zone of the germline is shown on Table S4.

**Quantification of PLK-2, HIM-8pT64, SUN-1pS12, and DSB-2 staining**

We used the presence of 2 or more aggregates of PLK-2, HIM-8 pT64, or SUN-1 pS12 on the nuclear envelope as markers of active chromosome movement. To determine the percentage of rows of nuclei displaying these markers we first counted the total number of rows of nuclei between the start of transition zone and the end pachytene. Then, starting from the beginning of transition zone, we identified the last row in which at least 50% of nuclei had 2 or more aggregates of PLK-2, HIM-8 pT64, or SUN-1 pS12 and counted the number of rows of nuclei included in that section of the germline. The number of rows scored as positive was then normalized to the total number of rows of nuclei in each germline. The same procedure was used to determine the region of DSB-2 positive staining, which we used as a marker of competence for DSB formation. Three germlines per genotype and condition were used for quantification. In Figure 7B, PLK-2 aggregates were quantified by dividing the region between meiosis onset and the end of pachytene into six equal-size regions and counting the number of nuclei with 0, 1, or >1 PLK-2 aggregates in each region.

**Western-blot**

Whole-worm protein extracts were prepared by picking 150 young adult worms of desired genotype into 1X Laemmli buffer and subjecting them to three cycles of freeze-thawing before boiling the samples for 5 minutes. Protein extracts were run on 10% acrylamide gels (Bio-Rad) and transferred onto nitrocellulose membrane for 1 hour at 4 $^{o}$C. Following 1 hour blocking in 1X TBS 0.1% Tween containing 5% dried milk, the nitrocellulose membranes were incubated overnight with rabbit anti-DSB-2 (Rosu et al., 2013) and goat anti-actin (Santa Cruz) primary antibodies, used at 1:1000 and 1:3000 respectively diluted in blocking buffer. After 3 washes of 10 minutes each with 1X TBS 0.1% Tween, membranes were incubated for one hour at room temperature with HRP-conjugated goat anti-rabbit (Jackson Immunoresearch) (1:5000) and HRP-conjugated donkey anti-goat IgG (Sigma) (1:8000) antibodies in blocking buffer. Finally, membranes were washed with 1x TBS 0.1% Tween (3 times for 10 minutes) and treated with ECL$^{TM}$ Western Blotting detection kit (GE Healthcare).

**RNA interference (RNAi)**

All RNAi experiments were carried out by feeding worms with HT115 bacteria transformed with a vector for IPTG-inducible expression of dsRNA. *syp-1* RNAi was performed using clone V-10P20 from the Ahringer library. Bacteria containing the *syp-1* vector, as well as empty vector (HT115) control, were both grown overnight at 37 °C in LB with 50 µg/ml ampicillin. Cultures were then centrifugated, collected and resuspended in 1 ml LB, before seeding 100 µl of this culture onto NGM agar plates containing 1 mM IPTG and 25 µg/ml ampicillin. When RNAi experiments were combined with auxin-mediated protein degradation RNAi plates also contained 1 or 4 mM auxin. Plates were incubated overnight at 37 °C to induce the expression of dsRNA. Eggs from the indicated strains were hatched on RNAi plates and maintained in these plates until they became young adults (18-24 hours post-L4) and were used for experiments.

**Viability screening**

L4 worms from the indicated strains were individually picked and transferred onto new plates every 12 hours, when the total number of embryos on each plate was counted. The presence of dead embryos (embryonic lethality) was assessed 24 hours after the mother had been removed from each plate. Five individual worms from each genotype were scored, and the total number of embryos counted is shown (n).

##### ***In-vivo*** **imaging**

Adult hermaphrodites were immobilized on a slide using a hydrogel-microbead solution similar to previously described protocol (Dong et al., 2018). First, adult worms treated for two hours in OP50 seeded plates containing 1mM auxin or no auxin controls were placed on a 3 µl droplet containing 0.5 mM levamisole dissolved in S medium to help immobilization. Next, 7 µl of precooled hydrogel-microbead solution (45% Pluronic F-127 (Sigma #P2443) mixed with 30 µm polystyrene microbeads (Sigma #84135) were added around the droplet and worms were overlaid with a coverslip containing also 7 µl of precooled hydrogel-microbead solution. Coverslips were sealed using nail polish and imaged immediately in a Delta Vision deconvolution system (Applied precision). 0.8 µm spaced Z-stack images were acquired with a time-lapse of 10 seconds over 7 minutes with the following settings: 50% power, 25 ms exposure, 60x microscope lens and 1024x1024 image size.

##### **RNAseq: RNA extraction**

Four biological replicates were used for each condition. For each replicate, fifty 18-24 hours post L4 worms were treated for four hours in OP50 seeded NG plates containing 4 mM auxin or to control plates without auxin. Next, adult worms were collected into 400 µl of M9 salt buffer in RNase-free tubes. After the worms settled to the bottom, the majority of the M9 was removed. 1 ml of Trizol (Thermo Fisher #15596026) and 100 µl of 0.5 mm of glass beads (Sigma #Z250465) were added to each tube and immediately frozen at -80 °C. Samples were thawed in ice and bead-beaten with a FastPrep-24 5G instrument (MP Biomedicals) with the following settings: speed 4 m/s, 3 cycles, and run time of 20 seconds. After the beads settled at the bottom of the tube, the solution was transferred to a new RNase-free tube. 200 µl of chloroform was added and tubes were vigorously mixed by hand for 15 sec. Samples were then centrifuged at 12000 g for 15 minutes at 4 °C and the resulting clear upper-phase was transferred to a new tube, where 500 µl of isopropanol and 1 µl of glycogen were added prior to storing the sample overnight at -20 °C. Next, samples were spun at 12000 g for 10 minutes at 4 °C, the supernatant was discarded and the pellet washed with 1 ml of fresh 75% ethanol. After 5 minutes centrifugation at 7500 g and 4 °C, ethanol was discarded, pellet briefly air-dried and resuspended with 15 µl of RNase-free water.

**RNAseq: sequencing and analysis**

Paired end 100bp libraries were sequenced on Illumina Hiseq 2500 and Raw basecall files were converted to fastq files using Illumina's bcl2fastq (version 2.1.7). Reads were aligned to the *Caenorhabditis elegans* genome (ce10) using Tophat2 version 2.0.11 (Kim et al., 2013) with default parameters. Mapped reads that fell on genes were counted using featureCounts from Rsubread package (Liao et al., 2019). Generated count data were then used to identify differentially expressed genes using DESeq2 (Love et al., 2014). Genes with very low read counts were excluded.

RNAseq data set GEO accession number: GSE134989 (private access token: yfopmksedpuhrwj)

| Genotype | Zone 1 | Zone 2 | Zone 3 | Zone 4 | Zone 5 | Zone 6 |
|---|---|---|---|---|---|---|
| *rec-8$^{3XTEV}$::GFP; rec-8* (-TEV) | 131 | 135 | 163 | 160 | 133 | 120 |
| *rec-8$^{3XTEV}$::GFP; rec-8* (+TEV) | 172 | 166 | 185 | 146 | 126 | 113 |
| *rec-8$^{3XTEV}$::GFP; rec-8; spo-11* (-TEV) | 114 | 124 | 154 | 144 | 133 | 115 |
| *rec-8$^{3XTEV}$::GFP; rec-8; spo-11* (+TEV) | 117 | 143 | 136 | 137 | 124 | 114 |
| *syp-2::AID* (-Auxin) | 241 | 165 | 215 | 169 | 139 | 172 |
| *syp-2::AID* (+Auxin) | 174 | 194 | 172 | 166 | 155 | 98 |

**Table S1. Number of nuclei analysed for RAD-51 staining in each region of the gonad in different genotypes**

**Table S2 (see excel file). Changes in gene expression of 335 meiotic genes following auxin-mediated depletion of SC component SYP-2 compared to untreated controls.**

Description of column values:

baseMean = average of the normalized count values for all samples (is a just the average of the normalized count values, dividing by size factors, taken over all samples)

log2FoldChange = effect size estimate (Treatment vs Control = how much the gene expression has changed due to Treatment). It's reported as log scale to base 2 (a value of 1.5 means 2^1.5 = 2.82 increase in expression)

lfcSE = uncertainty associated with log2FoldChange

stat = Wald statistic

pvalue = Wald test p-value. probability that observed log2FoldChange would be observed under null hypothesis

padj = BH adjusted p-values. Fraction of false positives given a gene's pvalue (method = Benjamini-Hochberg)

| Strain | Genotype |
|---|---|
| ATGSi355 | *fqSi16 II ; rec-8(ok978) IV* |
| ATGSi441 | *fqSi15 II ; coh-3(gk112) coh-4(1857) V* |
| ATG415 | *smc-1 (fq64[smc-1::AID::GFP]) I ; eSi38 IV* |
| ATGSi392 | *fqSi16 II ; rec-8(ok978) spo-11(ok79) IV / nT1 [qls51] (IV;V)* |
| ATG298 | *syp-2 (fq30[syp-2::AID]) V ; eSi38 IV* |
| ATG506 | *syp-3 (syb1022[mScarlet::syp-3]) I ; fqSi13 II ; syp-2 (fq30[syp-2::AID]) V ; eSi38 IV* |
| ATG330 | *chk-2 (fq41[chk-2::AID]) V ; eSi38 IV* |
| ATG387 | *chk-2 (fq41[chk-2::AID]) V ; fqSi16 II ; rec-8(ok978) eSi38 IV* |
| ATGSi525 | *fqSi15 II ; coh-3(gk112) coh-4(1857) V ; htp-1(gk174) IV / nT1 [unc-? (n754) let-? qls50] (IV;V)* |
| ATG302 | *rec-8(fq32[rec-8::TEV]) IV ; coh-3(fq35[coh-3::TEV]) coh-4 (1857) V* |
| ATGSi578 | *fqSi16 II ; rec-8(ok978) IV ; cosa-1(fq42[cosa-1::HA]) III* |
| ATGSi23 | *fqSi18 II ; rec-8(ok978) IV* |
| ATG169a | *fqSi19 I ; coh-3(gk112) coh-4(1857) V* |
| ATG554 | *syp-2 (fq30[syp-2::AID]) V ; cosa-1(fq42[cosa-1::HA]) III ; eSi38 IV* |
| ATG323 | *fqSi17 II ; rec-8(ok978) IV ; eSi38IV* |

**Table S3. List of *C. elegans* strains created in this study.**

| Transgene | Genotype | Origin |
|---|---|---|
| *fqSi16* | [*Prec-8 rec-8*$^{3XTEV}$*::GFP 3'UTR rec*-8; *cb-unc-119(+)*] | This study |
| *fqSi15* | [*Pcoh-3 coh-3*::$^{3XTEV}$*::mCherry 3'UTR coh-3*; *cb-unc-119(+)*] | This study |
| *fqSi13* | [*Pplk-2 plk-2*::GFP *3'UTR plk-2*; *cb-unc-119(+)*] | This study |
| *fqSi23* | [*Prec-8 rec-8::GFP 3'UTR rec*-8; *cb-unc-119(+)*] | *Crawley et al., 2016* |
| *fqSi18* | [*Pcoh-3 coh-3*::*mCherry 3'UTR coh-3*; *cb-unc-119(+)*] | This study |
| *fqSi17* | [*Prec-8 rec-8::AID::GFP 3'UTR rec*-8; *cb-unc-119(+)*] | This study |
| *ieSi38* | [*Psun-1 TIR1::mRuby 3'UTR sun-1; cb-unc-119(+)*] | Zhang *et al.*, 2015 |

**Table S4: Transgenes used in this study.**

| Alelle | sgRNA | Repair template |
|---|---|---|
| *rec-8(fq32[rec-8$^{3XTEV}$])* | CTGGGGTGCTGGTTGTTCCA | ttgtctctattgctctgctcccgagtgaaaccgtggagcaggagaatttgtatt<br>ttcagggtgcttctgaaaacctttacttccaaggagagctcgaa<br>aatctttatttccagggaccagcaccccaggagcctattcaagagcccattcaa |
| *Coh-3(fq35[coh-3$^{3XTEV}$])* | GGAACTTTGGGAATTCTTAT | ctcggaaaacatggatctagatgacgaagtttcgattgagaatttgtatt<br>ttcagggtgcttctgaaaacctttacttccaaggagagctcg<br>aaaatctttatttccagggaccgatcaggattcccaaagttcctcttagt<br>gctacaaagaaacaaa |
| *syp-2(fq30[syp-2::AID])* | ATTTATAACTTGTCAGCCCA | Left oligo:<br>aacttctggatttggttgaaacgctcgagccgtgggcagataaattg<br>atgcctaaagatccagccaaacctccggccaaggcacaagttgtgggatggccaccggtga<br>gatcataccggaagaac<br>Right oligo:<br>tgggatggccaccggtgagatcataccggaagaacgtgatggtttcctgccaaaaatcaagcg<br>gtggcccggaggcggcggcgttcgtgaagtaaatcatctgttgtattcaatttccttgttttattacat |
| *chk-2(fq41[chk-2::AID])* | AATGTGAACAACGTTCCACG | Left oligo:<br>aaaatttcaatttttttagccttaaaaaccctattttcaggcaaaaatgggaggtggagg<br>tggagctatgcctaaagatccagccaaacctccggccaaggcacaagttgtgggatggcca<br>ccggtgagatcataccggaagaac<br>Right oligo:<br>tgggatggccaccggtgagatcataccggaagaacgtgatggtttcctgccaaaaatcaagc<br>ggtggcccggaggcggcggcgttcgtgaagtagacaacgttccacgtgggtccccaccactt<br>caccgccgcaaaagtccg |
| *cosa-1(fq42[cosa-1::HA])* | TGTCAGAGATGGTAGTTACG | cagaatgagagtattccggaatgcagcacctcctcgtacccatacgatgtt<br>ccagattacgcttaactaccatctctgacagcacctctttgtcgccgat |
| *smc-1(fq64[smc-1::AID::GFP])* | TCCATAGCTACAATGAGTAA | Left oligo:<br>Tgactgaaaatcctcctactccttccatagctacaatgccaaaagatccagcc<br>Aaacctccggccaaggcacaagttgtgggatggccaccggtgagatcataccggaagaac<br><br>Right oligo:<br>tgggatggccaccggtgagatcataccggaagaacgtgatggtttcctgccaaaaatcaag<br>cggtggcccggaggcggcggcgttcgtgaagatgTCtaaaggagaagaacttttcactggagttgtcccaa |

**Table S5. CRISPR alleles created in this study.**

### Figure S1

A

| Genotype | Embryonic lethality (%) | n |
|---|---|---|
| WT (N2) | 2.32 | 1505 |
| *REC-8::GFP; rec-8 (ok978)* | 1.6 | 1664 |
| *REC-8$^{3XTEV}$::GFP; rec-8 (ok978)* | 2.44 | 1501 |
| *rec-8 (ok978)* | 86.89 | 929 |
| *COH-3::mcherry; coh-3 (gk112); coh-4 (tm1857)* | 2.41 | 1412 |
| *COH-3$^{3XTEV}$::mcherry; coh-3 (gk112); coh-4 (tm1857)* | 2.04 | 1306 |
| *coh-3 (gk112); coh-4 (tm1857)* | 97.57 | 774 |

B

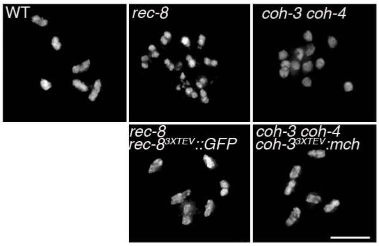


C

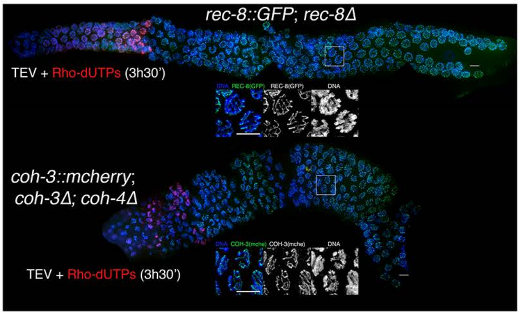


D

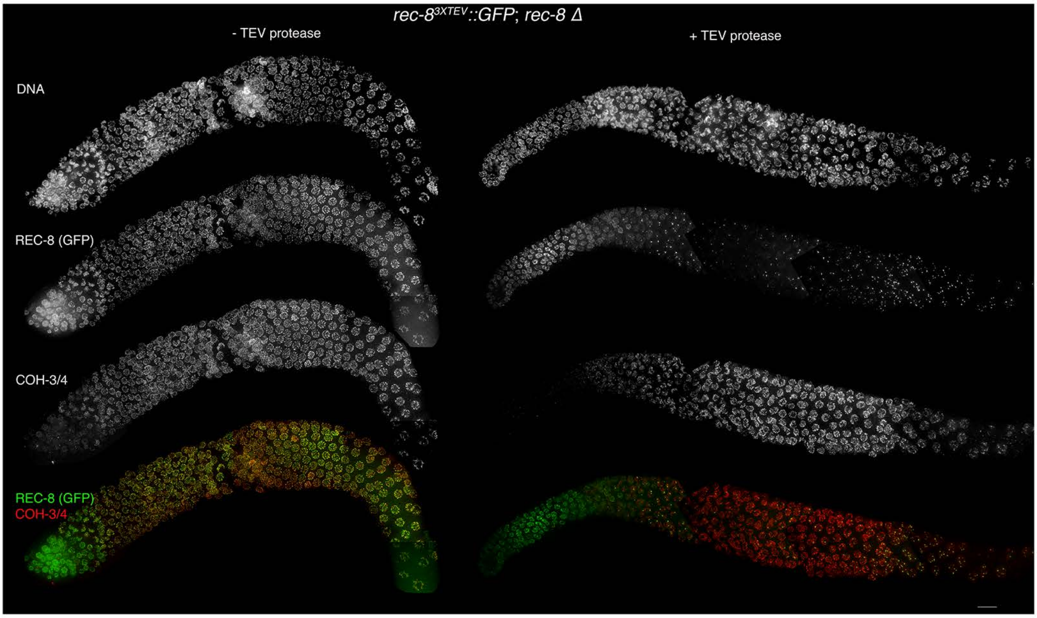


E

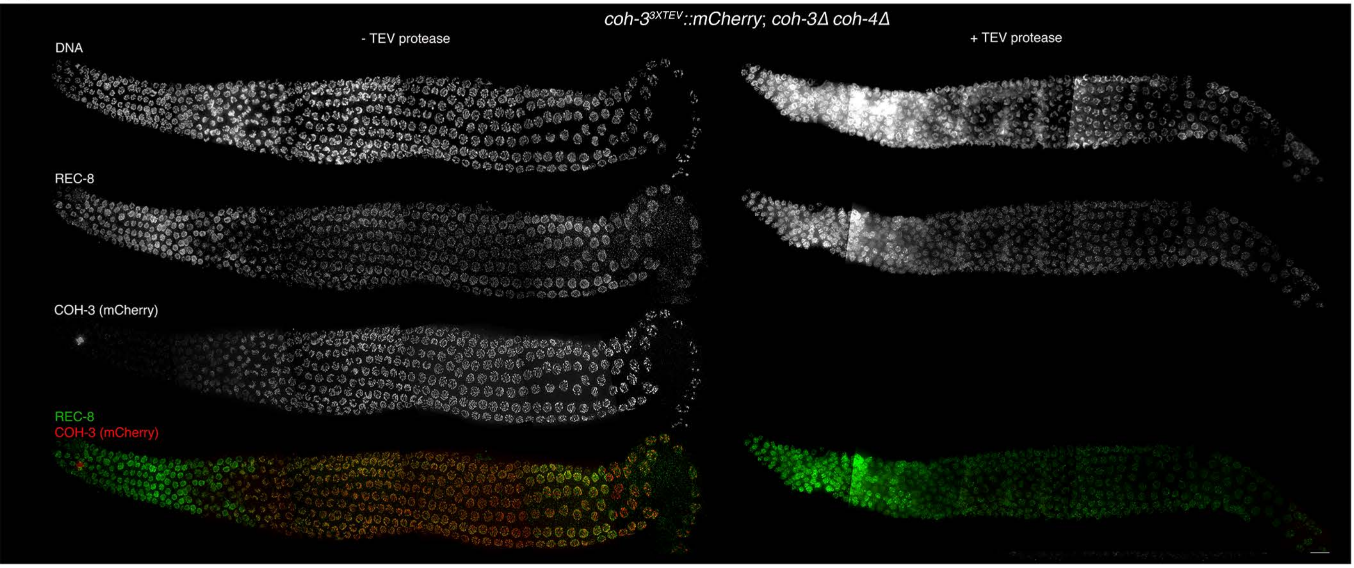

**Figure S2**

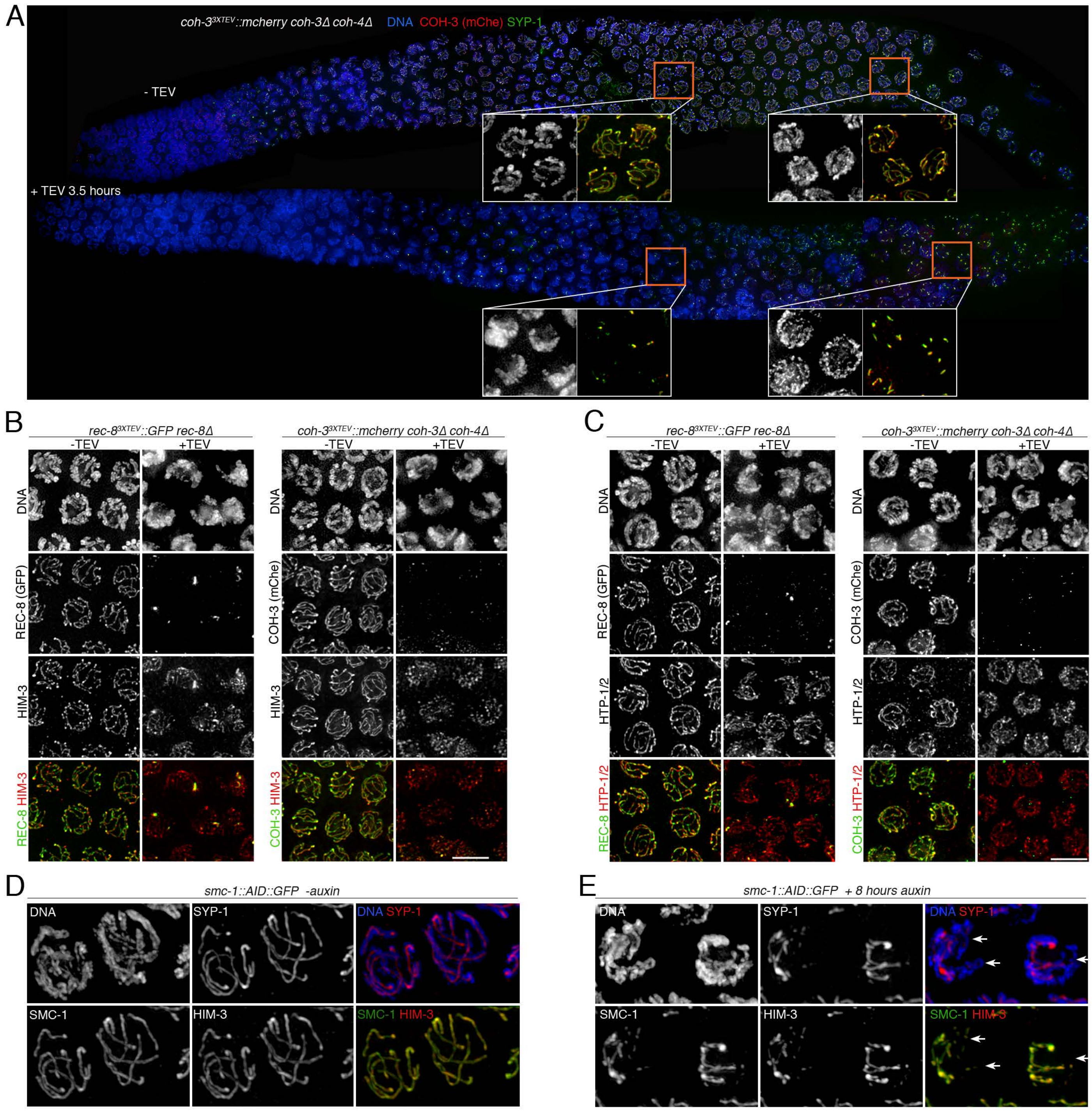

A
coh-3^3XTEV::mcherry coh-3Δ coh-4Δ DNA COH-3 (mChe) SYP-1
- TEV
+ TEV 3.5 hours
B
rec-8^3XTEV::GFP rec-8Δ
coh-3^3XTEV::mcherry coh-3Δ coh-4Δ
-TEV
+TEV
DNA
REC-8 (GFP)
COH-3 (mChe)
HIM-3
REC-8 HIM-3
COH-3 HIM-3
C
rec-8^3XTEV::GFP rec-8Δ
coh-3^3XTEV::mcherry coh-3Δ coh-4Δ
HTP-1/2
REC-8 HTP-1/2
COH-3 HTP-1/2
D
smc-1::AID::GFP -auxin
DNA
SYP-1
DNA SYP-1
SMC-1
HIM-3
SMC-1 HIM-3
E
smc-1::AID::GFP + 8 hours auxin

### Figure S3

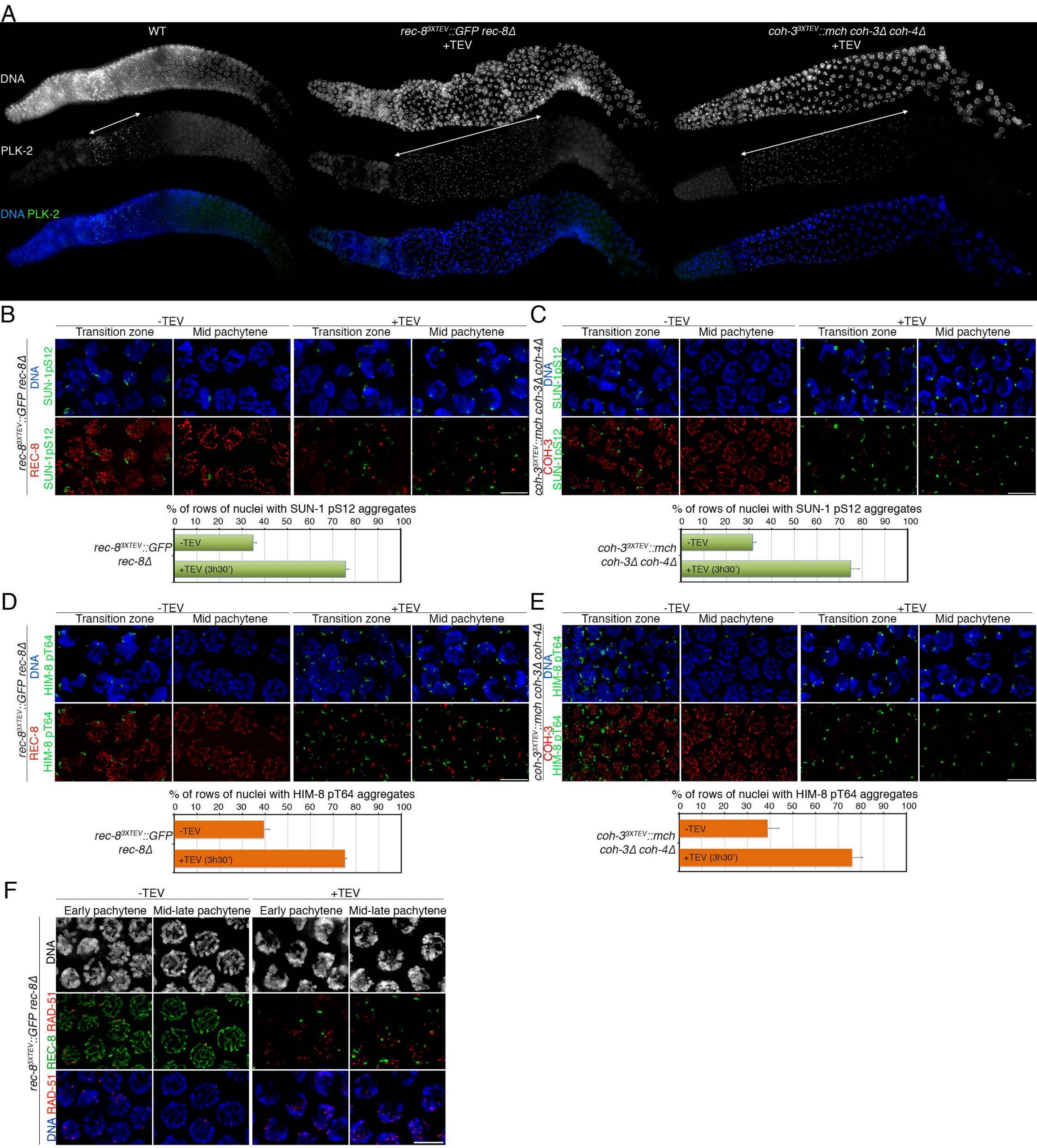
A
WT
rec-8^3XTEV::GFP rec-8Δ +TEV
coh-3^3XTEV::mch coh-3Δ coh-4Δ +TEV
DNA
PLK-2
DNA PLK-2
B
-TEV
+TEV
Transition zone
Mid pachytene
rec-8^3XTEV::GFP rec-8Δ
DNA SUN-1pS12
REC-8 SUN-1pS12
% of rows of nuclei with SUN-1 pS12 aggregates
0 10 20 30 40 50 60 70 80 90 100
rec-8^3XTEV::GFP rec-8Δ
-TEV
+TEV (3h30')
C
coh-3^3XTEV::mch coh-3Δ coh-4Δ
DNA SUN-1pS12
COH-3 SUN-1pS12
% of rows of nuclei with SUN-1 pS12 aggregates
coh-3^3XTEV::mch coh-3Δ coh-4Δ
D
DNA HIM-8 pT64
REC-8 HIM-8 pT64
% of rows of nuclei with HIM-8 pT64 aggregates
E
DNA HIM-8 pT64
COH-3 HIM-8 pT64
% of rows of nuclei with HIM-8 pT64 aggregates
F
Early pachytene
Mid-late pachytene
DNA
REC-8 RAD-51
DNA RAD-51

### Figure S4

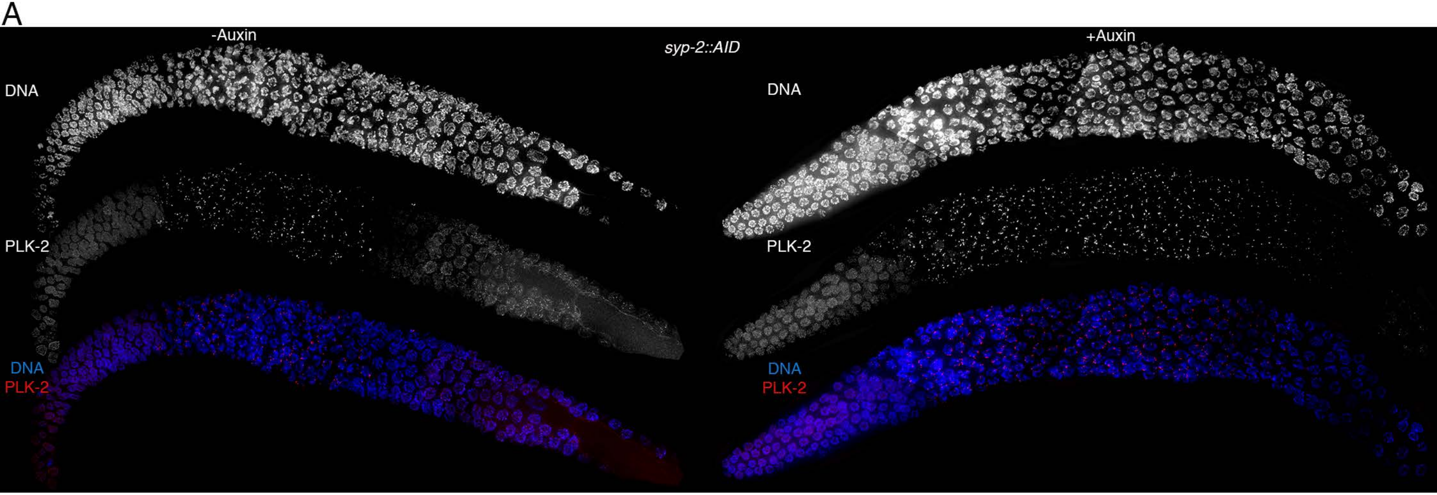

A
-Auxin
syp-2::AID
+Auxin
DNA
PLK-2
DNA
PLK-2
DNA
PLK-2
DNA
PLK-2


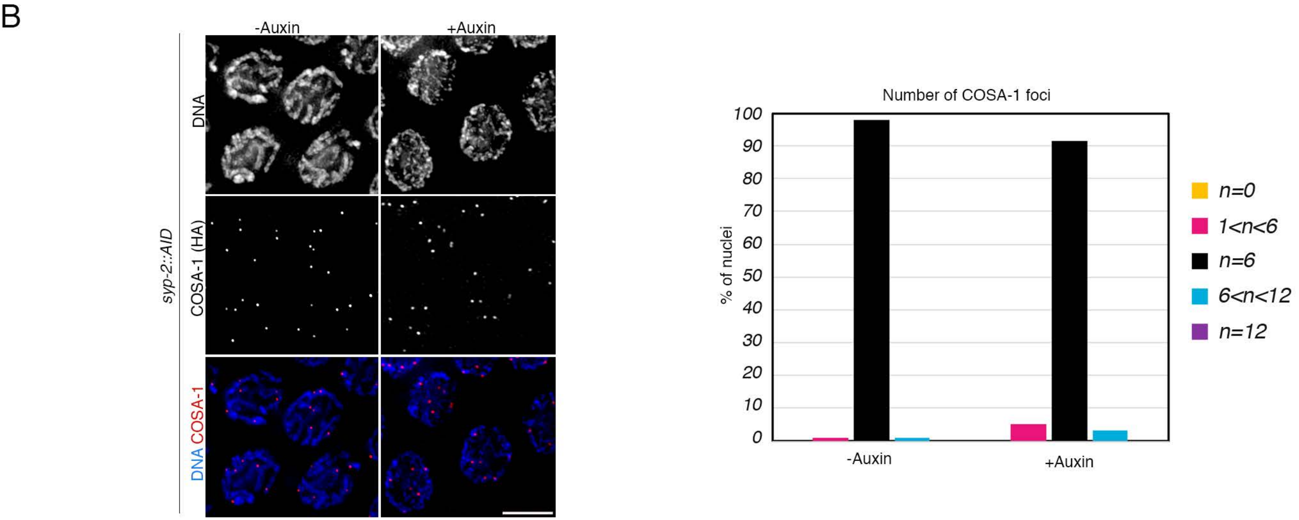

B
-Auxin
+Auxin
syp-2::AID
DNA
COSA-1 (HA)
DNA COSA-1
Number of COSA-1 foci
% of nuclei
100
90
80
70
60
50
40
30
20
10
0
-Auxin
+Auxin
n=0
1<n<6
n=6
6<n<12
n=12

#### Figure S5

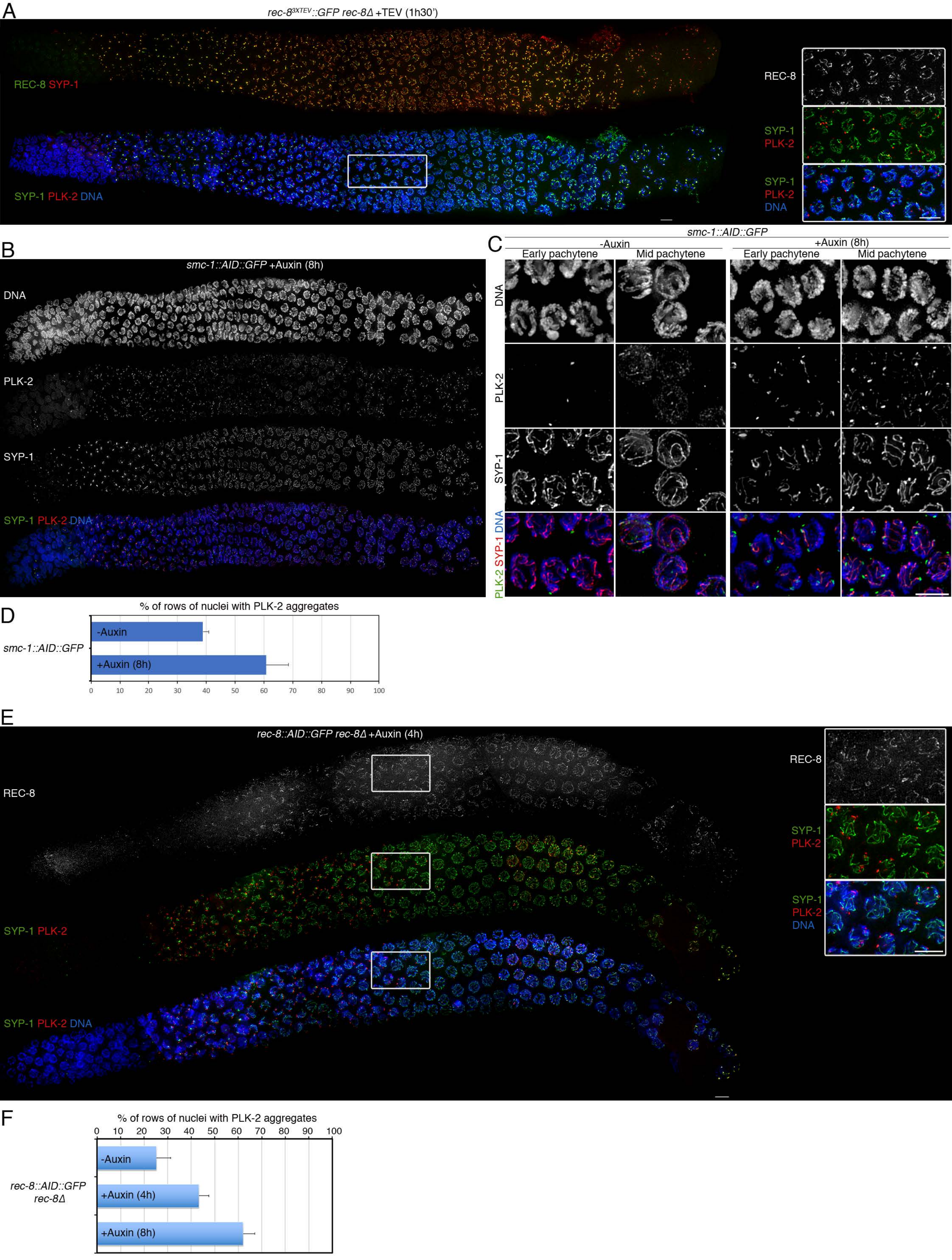

A
rec-8^3XTEV::GFP rec-8Δ +TEV (1h30')
REC-8 SYP-1
SYP-1 PLK-2 DNA
REC-8
SYP-1
PLK-2
SYP-1
PLK-2
DNA
B
smc-1::AID::GFP +Auxin (8h)
DNA
PLK-2
SYP-1
SYP-1 PLK-2 DNA
C
smc-1::AID::GFP
-Auxin
+Auxin (8h)
Early pachytene
Mid pachytene
Early pachytene
Mid pachytene
DNA
PLK-2
SYP-1
PLK-2 SYP-1 DNA
D
% of rows of nuclei with PLK-2 aggregates
smc-1::AID::GFP
-Auxin
+Auxin (8h)
0 10 20 30 40 50 60 70 80 90 100
E
rec-8::AID::GFP rec-8Δ +Auxin (4h)
REC-8
SYP-1 PLK-2
SYP-1 PLK-2 DNA
REC-8
SYP-1
PLK-2
SYP-1
PLK-2
DNA
F
% of rows of nuclei with PLK-2 aggregates
0 10 20 30 40 50 60 70 80 90 100
rec-8::AID::GFP
rec-8Δ
-Auxin
+Auxin (4h)
+Auxin (8h)

#### Figure S6

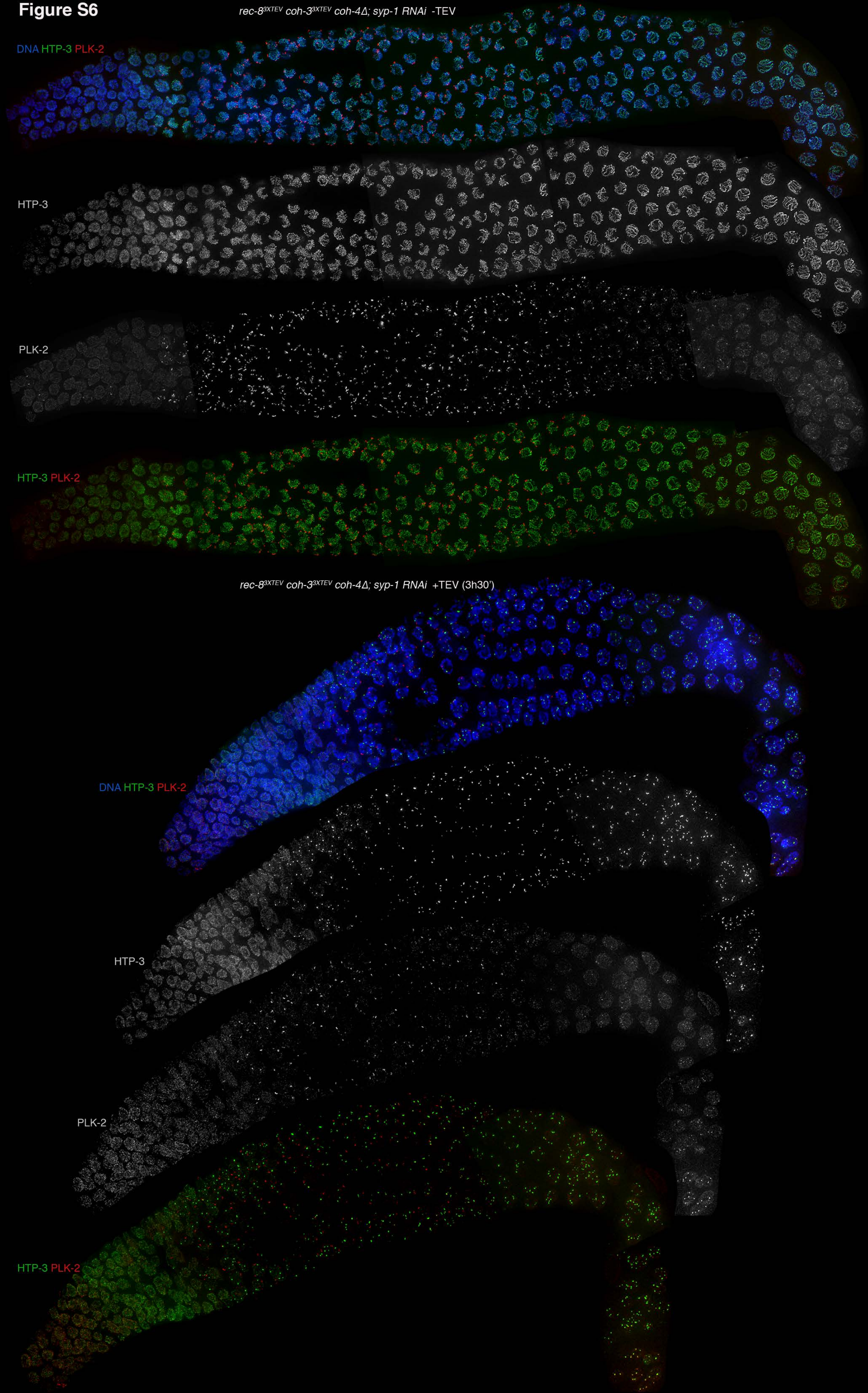

rec-8^3XTEV coh-3^3XTEV coh-4Δ; syp-1 RNAi -TEV
DNA HTP-3 PLK-2
HTP-3
PLK-2
HTP-3 PLK-2
rec-8^3XTEV coh-3^3XTEV coh-4Δ; syp-1 RNAi +TEV (3h30')
DNA HTP-3 PLK-2
HTP-3
PLK-2
HTP-3 PLK-2

**Figure S7**

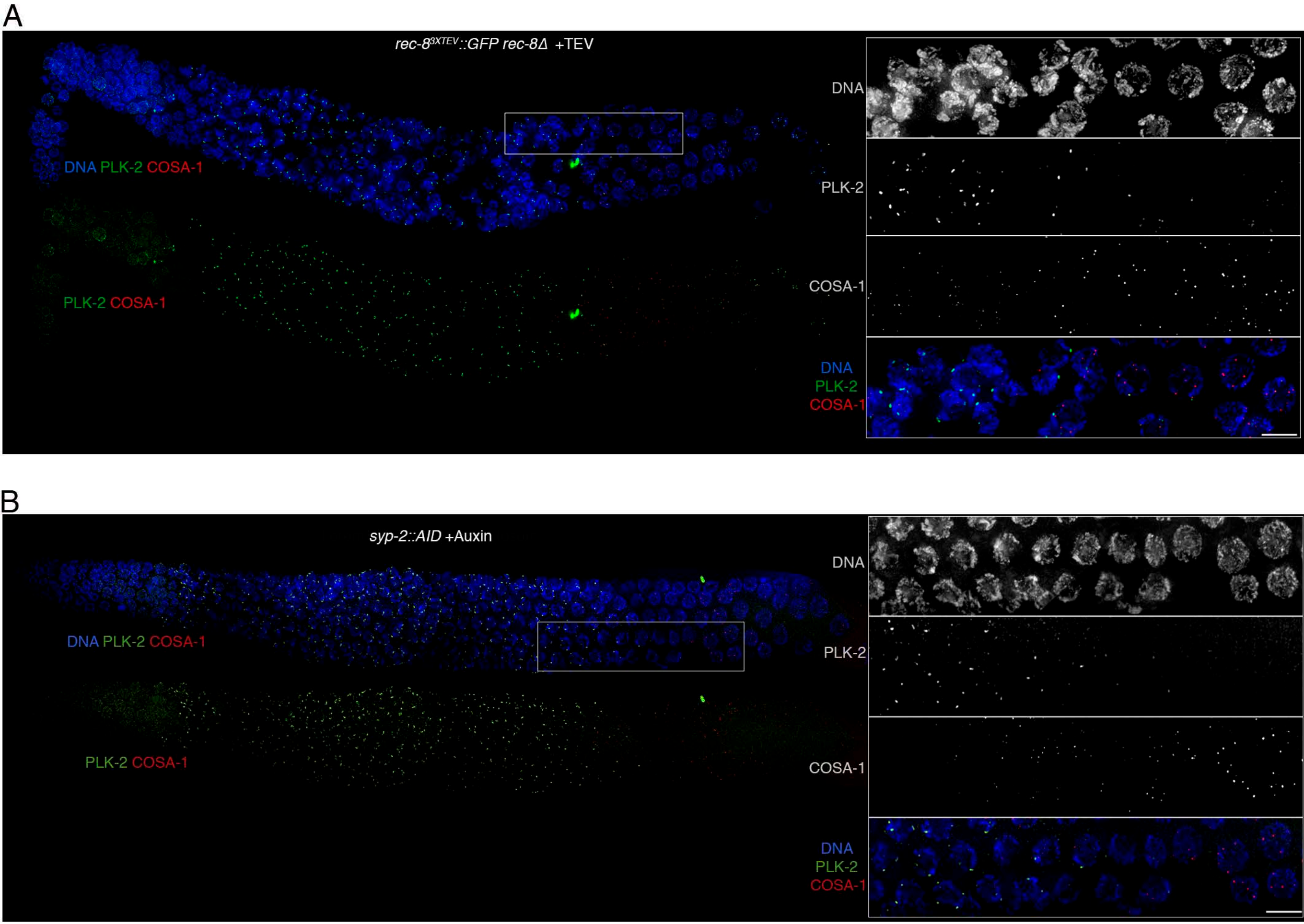
A
rec-8^3XTEV::GFP rec-8Δ +TEV
DNA PLK-2 COSA-1
PLK-2 COSA-1
DNA
PLK-2
COSA-1
DNA
PLK-2
COSA-1
B
syp-2::AID +Auxin
DNA PLK-2 COSA-1
PLK-2 COSA-1
DNA
PLK-2
COSA-1
DNA
PLK-2
COSA-1
